# Sec23IP recruits the Cohen syndrome factor VPS13B/COH1 to ER exit site-Golgi interface for tubular ERGIC formation

**DOI:** 10.1101/2024.02.28.582656

**Authors:** Yuanjiao Du, Xinyu Fan, Chunyu Song, Juan Xiong, Wei-Ke Ji

**Author notes:** Equal contribution (Yuanjiao Du, Xinyu Fan).

## Abstract

VPS13B/COH1 is the only known causative factor for Cohen syndrome, an early-onset autosomal recessive developmental disorder with intellectual inability, developmental delay, joint hypermobility, myopia and facial dysmorphism as common features, but the molecular basis of VPS13B/COH1 in pathogenesis is unknown. Here, we identify Sec23 interacting protein (Sec23IP) at ER exit site (ERES) as a VPS13B adaptor that recruits VPS13B to ERES-Golgi interfaces. VPS13B interacts directly with Sec23IP via the VPS13 adaptor binding domain (VAB), and the interaction promotes the association between ERES and the Golgi. Disease-associated missense mutations of VPS13B-VAB impair the interaction with Sec23IP. Knockout of VPS13B or Sec23IP blocks the formation of tubular ERGIC, an unconventional cargo carrier that expedites ER-to-Golgi transport. In addition, depletion of VPS13B or Sec23IP delays ER export of procollagen, suggesting a link between procollagen secretion and joint laxity in patients with Cohen disease. Together, our study reveals a crucial role of VPS13B-Sec23IP interaction at ERES-Golgi interface in the pathogenesis of Cohen syndrome.

## Introduction

The early secretory pathway, which consists of vesicular traffic between the endoplasmic reticulum (ER) and the Golgi apparatus, occurs constitutively in mammalian cells. The pathway is crucial for the constant supply of secretory and plasma membrane lipids/proteins and is considered essential for general cell function and survival^1^. Neurons exhibit a high intensity of membrane dynamics and protein/lipid transport, with differential and polarized transport towards the somato-dendritic and axonal plasma membrane domains^2^. Mutations in genes encoding components of the early secretory pathway are known to cause neurological or developmental disorders that manifest early in life^3^. These rare disorders are associated with autosomal recessive mutations in coat proteins, membrane tethering and fusion complexes^4^, such as subunits of coat protein complex I and II (COPI and COPII)^5, 6^, subunits of the transport protein particle complex (TRAPP)^7^, members of the YIP1 domain family (YIPF)^8^ and a member of the SNAP receptor family (SNARE)^9^.

Cohen syndrome (MIM 216550), a rare recessive developmental disorder was first described by Cohen and co-workers^10, 11^, with a variable clinical presentation characterized mainly by developmental delay, mental retardation, joint laxity, microcephaly, typical facial dysmorphism, progressive pigmentary retinopathy, severe myopia and intermittent neutropenia. Among these features, developmental delay, early onset myopia, joint laxity and facial dysmorphism were the cardinal clinical phenotypes present in all patients with Cohen syndrome^12^.

All patients with Cohen disease were homozygous or compound heterozygous for mutations in a gene encoding vacuolar protein sorting-associated protein 13B (VPS13B, also known as COH1)^12^. VPS13B is a member of the bridge-like repeating β-groove (RBG) lipid transfer protein family^13^. The human genome contains four VPS13 genes (VPS13A, VPS13B, VPS13C and VPS13D genes)^14^. Each of them is directly associated with certain human diseases, and therefore these proteins are of great biomedical interest^15^. Previous studies have shown that VPS13A and VPS13C are lipid transporters at ER-associated membrane contact sites (MCSs), including ER-mitochondria/plasma membrane (VPS13A) and ER-late endosome/lysosome MCSs (VPS13C)^16–18^. VPS13D has been reported to play a role in mitophagy in Drosophila^19, 20^ and is localized at ER-mitochondrial/peroxisomal contacts^21^, and promotes peroxisome biogenesis^22^. Our previous results have shown that VPS13D plays a regulatory role in ER-mitochondrial MCSs^22^ and facilitates LD remodeling under starvation^23^. To date, the relationship between VPS13B and MCSs is unknown.

VPS13B has been involved in several cellular processes, such as Golgi integrity and neurite outgrowth^24, 25^, cargo recycling transport^26^, acrosome biogenesis^27^ and LD dynamics^28^. However, the molecular basis underlying these cellular functions and the link between VPS13B and the pathogenesis of Cohen syndrome are still lacking. In this study, we sought to explore the molecular mechanism of these cellular functions mediated by VPS13B with the aim of providing mechanistic insight into the pathology of Cohen syndrome.

## Results

### VPS13B localized to the Golgi via binding to PI4P

First, we investigated the cellular localization of endogenous VPS13B using immunofluorescence (IF). The endogenous VPS13B was colocalized with GM130, a cis-/medial Golgi marker (Fig. 1A). The VPS13B fluorescence was completely lost in CRISPR-Cas9-mediated VPS13B knockout HeLa cells (VPS13B KO; Fig.S1A, B), confirming the specificity of the VPS13B antibody utilized in IF. Consistently, GFP tagged VPS13B (VPS13B-GFP) was well co-localized with the Golgi (Fig. S1C).

**Fig. 1.**
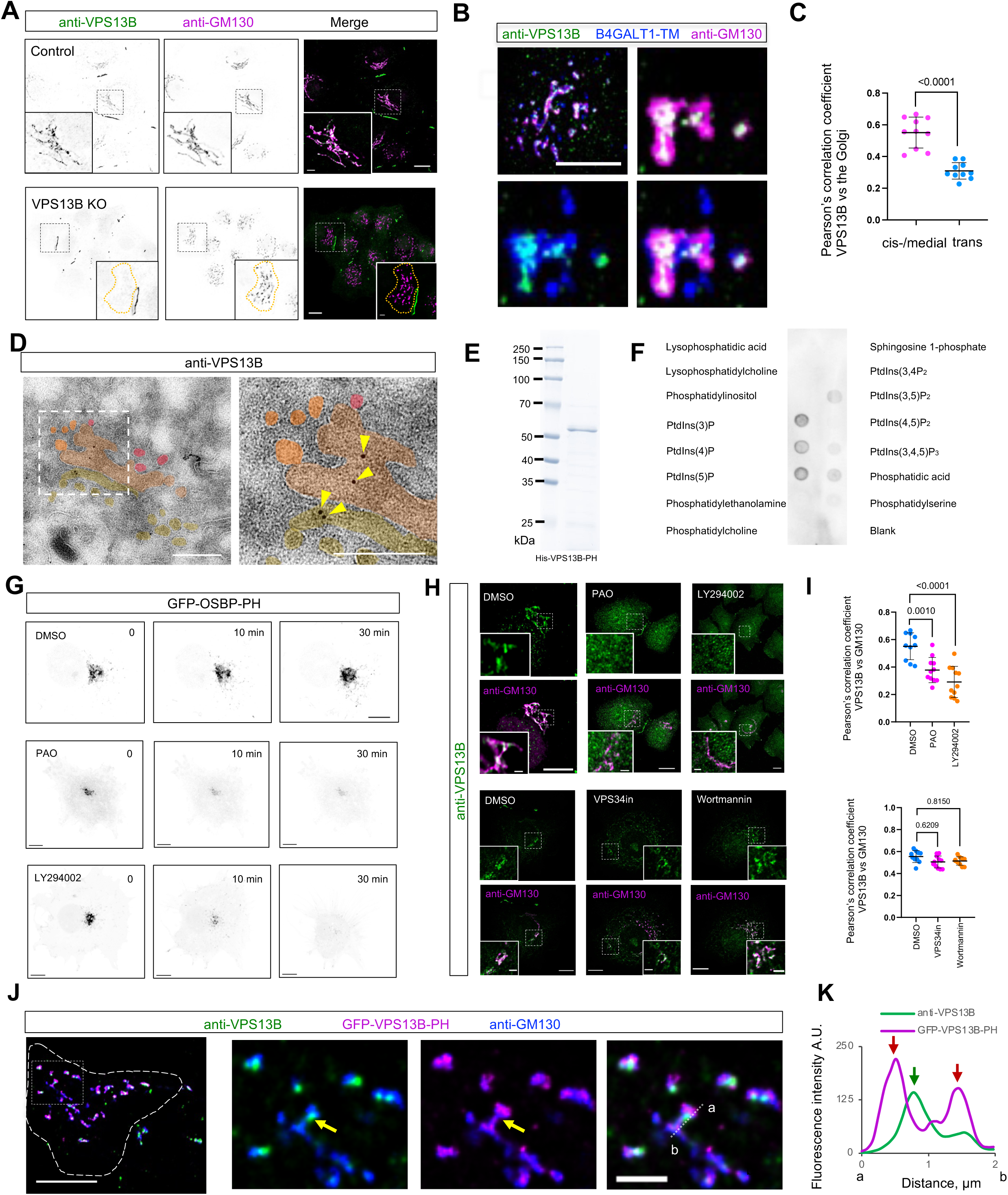
VPS13B preferentially localizes to the cis-Golgi via binding to PI4P. (A) Representative images of a fixed control (top) or VPS13B KO HeLa cell (bottom) stained with VPS13B antibody (green) and GM130 antibody (magenta) with insets. (B) Representative images of a fixed HeLa cell transiently expressing GFP-B4GALT1-TM (blue) stained with VPS13B antibody (green) and GM130 antibody (magenta) with an inset. (C) Pearson’s correlation coefficient of endogenous VPS13B vs the cis-/medial Golgi (GM130; 10 cells) or the trans-Golgi (B4GALT1-TM; 10 cells) in 3 independent experiments. Mean ± SD. Two tailed unpaired student t test. (D) Representative EM micrograph of a cryosection of a fixed HeLa cell with endogenous VPS13B labeled by immunogold beads with an inset shown on the right. Yellow arrowheads denote VPS13B signals on the cis-/medial Golgi. (E) Coomassie blue staining of purified His tagged VPS13B PH-like domain. (F) The PIP Strip assays using purified His-VPS13B-PH as in (**E**). (G) Representative images of live HeLa cells transiently expressing a PI4P probe (GFP-OSBP-PH) upon control (DMSO; top), PAO (middle; 10 μM) and LY294002 (bottom; 300 μM) treatments with three timepoints (0, 10 min, 30 min). (H) Representative images of a fixed HeLa cell stained with VPS13B antibody (green) and GM130 antibody (magenta) treated with control (DMSO; top), PAO (10 μM; 30 min), LY294002 (300 μM; 30 min), Wortmannin (10 μM; 6 h) and VPS34in (1 μM; 6 h) with insets. (I) Pearson’s correlation coefficient of endogenous VPS13B vs the Golgi upon treated with DMSO (10 cells), PAO (11 cells), LY294002 (11 cells), Wortmannin (11 cells) and VPS34in (10 cells) in 3 independent experiments as in (**I**). Ordinary one-way ANOVA with Tukey’s multiple comparisons test. Mean ± SD. (J) Representative images of fixed HeLa cells transiently expressing GFP-VPS13B-PH (magenta) stained by VPS13B antibody (green) and GM130 antibody (blue) with insets on the right. Yellow arrowheads denote a punctae of endogenous VPS13B on the Golgi. (K) Line-scan analysis for (**J**) with red arrows indicating GFP-VPS13B-PH foci while a green arrow denoting anti-VPS13B foci on a Golgi ribbon. Scale bar, 10 μm in the whole cell images and 2μm in the insets in (A, B, G, H & J). 2 μm in (D).

The Golgi apparatus is a highly polarized organelle consisting of cis-/medial, trans- and trans-Golgi network. We further investigated which compartment of the Golgi VPS13B was mainly localized. We used transmembrane domain of beta-1,4-galactosyltransferase 4 tagged by GFP (GFP-B4GALT1-TM) as a trans-Golgi marker. Endogenous VPS13B was better colocalized with GM130 than B4GALT1-TM (Fig. 1B), as shown by colocalization analyses (Fig. 1C), suggesting that VPS13B is preferentially localized to the cis-/medial Golgi.

To confirm this hypothesis, we performed ultrastructural studies using immuno-electron microscopy (EM). While the level of endogenous VPS13B was very low, we could still observe that VPS13B signals (yellow arrowheads) were present on the cis- or medial Golgi, but not on the trans-Golgi network (TGN) (Fig. 1D).

We next asked how VPS13B was associated with the cis-/medial Golgi. VPS13B contains a lipid transfer domain along its entire length, with a VPS13 adaptor binding domain (VAB), an ATG2-C domain and a PH-like domain at the C-terminus (CT)^29, 30^. We hypothesized that the PH-like domain of VPS13B might be responsible for targeting VPS13B to the Golgi by binding to phosphatidylinositol 4-phosphate (PI4P) on Golgi membranes. To test this hypothesis, we purified the PH-like domain (Fig. 1E) and investigated whether it bound to PI4P. In vitro PIPs strip assays showed that the purified VPS13B-PH bound to PI3P, PI4P, and PI5P (Fig. 1F), partially consistent with a previous study using cell lysate containing overexpressed VPS13B-GFP^26^.

Next, we investigated whether PI4P is required for VPS13B localization. Treatment with phenylarsine oxide (PAO), a PI4K inhibitor, significantly decreased PI4P levels in the Golgi, as indicated by a substantial reduction in florescence intensity of a PI4P probe (PH domain of oxysterol-binding protein) (Fig. 1G). In addition, LY249002, a PI3K inhibitor, could also reduce the PI4P level when used at high concentrations^31^. Therefore, we investigated whether the inhibition of PI4P synthesis by these two inhibitors would block the association of endogenous VPS13B with the Golgi. Indeed, the association was greatly reduced after the treatment with PAO or LY294002 (Fig. 1H, I). However, two PI3K inhibitors, wortmannin and VPS34in, were unable to reduce the association (Fig. 1H, I), suggesting that PI4P, but not PI3P, is responsible for the targeting of VPS13B to the Golgi.

Moreover, the PH domain of VPS13B alone was able to target the Golgi (Fig. 1J). Both endogenous full length (FL) VPS13B and the PH domain were localized on the cis-/medial Golgi, but appeared to be mutually excluded, as shown by line-scan analyzes (Fig. 1K). This suggests that FL-VPS13B and the PH domain may compete with each other for binding PI4P. Accordingly, VPS13B with a deletion of the PH domain lost the ability to target the Golgi (Fig. S1D), confirming the role of the PH domain in the targeting of VPS13B to the Golgi. Noteworthy, we speculated that PI4P might not be the only determinant factor for VPS13B localization in the cis-/medial Golgi because PI4P was also enriched in the trans Golgi or TGN.

The PH domain of VPS13B was shown to interact with Rab6^24^, a small GTPase residing on the Golgi. We thus investigated whether Rab6 was required for VPS13B localization on the Golgi. We found that suppression of Rab6A or Rab2A, another Golgi-resident small GTPase, with small interfering RNAs (siRNAs) did not inhibit the association of GFP-VPS13B-PH with the Golgi (Fig. S1E-G). Accordingly, overexpression of dominant-negative mutants of Rab6A (Rab6A-T27N) appeared not significantly reduce colocalization (Fig. S1H). Consistent with this, Rab6 activity was not required for the association of endogenous VPS13B with the Golgi, as depletion of Rab6A (Fig. S1I) or overexpression of Rab6A-T27N (Fig. S1J) did not strongly reduce colocalization between VPS13B and GM130 (Fig. S1K). Taken together, these results suggested that the PH domain was responsible for VPS13B targeting to the Golgi by binding to PI4P independent of Rab6.

### VPS13B did not stably associate with the ER via VAPs

Bridge lipid transporters are thought to function at ER-associated contacts, with a FFAT or phospho-FFAT motif at the NT recognizing the ER in a VAP-dependent manner^29^. Therefore, we investigated whether VPS13B associated with the ER via this mechanism. We have previously shown that FFAT motifs at the NT of VPS13B (residues 1-1500) were not sufficient for the association with the ER^28^. Next, we tested whether VPS13B targeted the ER via a phospho-FFAT motif^32^. A phospho-FFAT motif was found at the NT of VPS13B (551-GSTNQQDFSSGKSEDLGTV; Fig. S2A). The structure and position of this phospho-FFAT motif was similar to its paralog VPS13D^21^ (Fig. S2B). Co-immunoprecipitation (coIP) assays showed that the VPS13B-NT interacted with VAPB but not VAPA or MOSPD3 (Fig. S2C). A phosphomimetic S560D mutation moderately increases the interaction between VPS13B-NT and VAPB. However, our imaging data showed that neither the VPS13B-NT nor phosphomimetic mutants (S560D, S1403D and S1433D) was able to target the ER (Fig. S2D-G). These results suggest that VPS13B may not stably associate with the ER via binding to the known ER adaptors. To investigate whether VPS13B targets the ER via unknown adaptors, we performed mass spectrometry (MS) to identify proteins interacting with VPS13B on the ER.

### VPS13B interacted with Sec23IP at ERES-Golgi interface

Using co-immunoprecipitation (co-IP) followed by mass spectrometry, we identified a protein called Sec23 interacting protein (Sec23IP/p125) as a protein interacting with VPS13B (Fig. 2A). Interestingly, both VPS13B and Sec23IP are present in vertebrates and have no homolog in other organisms, such as yeast and *Caenorhabditis elegan*. The direct link between Sec23IP and diseases was currently lacking, but an orthologous Sec23IP gene in frogs was involved in the development of neural crest cells. Intriguingly, both VPS13B and Sec23IP were reported to be crucial in acrosome biogenesis during spermiogenesis in mice^33^. Importantly, CRISPR-Cas9-mediated KO of Sec23IP (Fig. S3A, B) also resulted in fragmentation of the Golgi that phenocopied the KO of VPS13B (Fig.S3C)^24,28^. These results strongly suggested a functional link between VPS13B and Sec23IP.

**Fig. 2.**
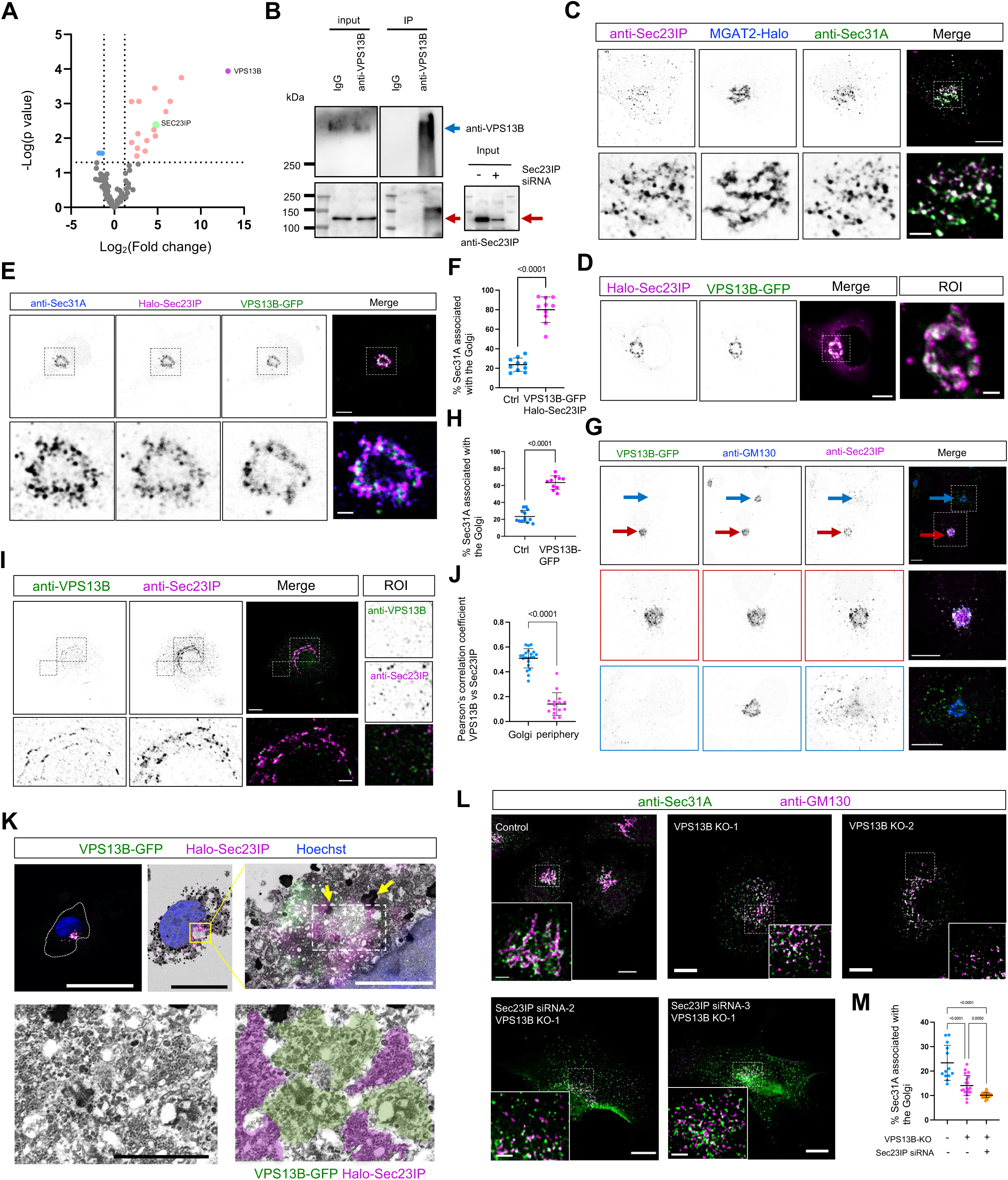
VPS13B interacts with Sec23IP at ERES-Golgi interfaces. (A) Volcano plot of protein candidates coIPed with VPS13B-GFP in HEK293 cells compared with protein candidates coIPed with GFP tag only. After removal of proteins that coIPed with GFP tag, candidates that were considered significant (−log [P value] > 1.3; P < 0.05) were labeled in orange (Log2 [fold change] > 0; increased in abundance) or blue (Log2 [fold change] < 0; decreased in abundance). (B) coIP assay showed an interaction between endogenous VPS13B and endogenous Sec23IP in HEK293 cells. Blue arrows denoted VPS13B while red arrows indicating Sec23IP in the blots. (C) Representative images of a fixed HeLa cell expressing MGAT2-Halo (blue; a cis-/medial Golgi marker) stained with Sec23IP antibody (magenta) and Sec31A antibody (green) with two insets. (D) Representative images of a live HeLa cell co-expressing Halo-Sec23IP (magenta) and VPS13B-GFP (green) with an inset to the right. (E) Representative images of a fixed HeLa cell co-expressing Halo-Sec23IP (magenta) and VPS13B-GFP (green) stained with Sec31A antibody (blue) with an inset on the bottom. (F) Percentage of Sec23IP puncta associated with the Golgi in either control (13 cells) or cells co-expressing Halo-Sec23IP and VPS13B-GFP (10 cells) as in (**D**). Mean ± SD. Two tailed unpaired student t test. (G) Representative images of fixed HeLa cells expressing VPS13B-GFP (green) stained with GM130 antibody (blue) and Sec23IP antibody (magenta) with two insets on bottom. Red arrows indicated a cell with the expression of VPS13B-GFP while blue arrows denoted a cell without VPS13B-GFP expression. (H) Percentage of Sec23IP puncta associated with the Golgi in either control (13 cells) or cells expressing VPS13B-GFP as in (H). Mean ± SD. Two tailed unpaired student t test. (I) Representative images of a fixed HeLa cell stained with VPS13B antibody (green) and Sec23IP antibody (magenta). One inset from the Golgi region was shown on the bottom while the other inset was shown to the right. (J) Pearson’s correlation coefficient of endogenous VPS13B vs endogenous Sec23IP at the Golgi (21 cells) or cell periphery (18 cells) in 3 independent experiments. Mean ± SD. Two tailed unpaired student t test. (K) HeLa cells expressing VPS13B-GFP, Halo-Sec23IP, and stained by DAPI, were fixed and imaged by 3D microscopy (top left panel) and processed for TEM. Fluorescence and TEM images were correlated (top middle panel) with insets showing associations between the cis-medial Golgi marked by VPS13B-GFP (green) and ERES labeled by Halo-Sec23IP (magenta). (L) Representative images of fixed control, VPS13B KO, or siRNA-mediated Sec23IP knockdown in VPS13B KO HeLa cells stained with GM130 antibody (magenta) and Sec31A antibody (green) with insets. (M) Percentage of Sec23IP puncta associated with the Golgi in either control (13 cells), VPS13B KO (20 cells) or VPS13B/Sec23IP double depleted cells (10 cells) as in (**L**). Mean ± SD. Ordinary one-way ANOVA with Tukey’s multiple comparisons test. Scale bar, 10 μm in the whole cell images and 2 μm in the insets in C, D, E, G, I and L; 0.2 μm in the insets in K.

To confirm that VPS13B interacts with Sec23IP at the endogenous levels, we performed co-IP assays. Endogenous Sec23IP could be co-pelleted by endogenous VPS13B (Fig.2B).

Next, we examined whether the interaction between Sec23IP and VPS13B was strong enough to mediate recruitment. Consistent with previous studies^34, 35^, both endogenous (Fig.2C) and exogenous Sec23IP (Fig. S3D) formed puncta over the cytosol that colocalized with Sec31A, an ER exit site (ERES) marker, with a substantial fraction of Sec23IP puncta tightly associated with the cis-/medial Golgi. Strikingly, a much larger proportion of Halo-Sec23IP puncta was recruited to the cis-/medial Golgi when VPS13B-GFP was co-expressed (Fig.2D). This suggests that ERES is strongly recruited to the Golgi via VPS13B-Sec23IP interaction.

To test this hypothesis, we examined the spatial relationship between ERES (anti-Sec31A) and the cis-/medial Golgi upon co-expression of VPS13B-GFP and Halo-Sec23IP. IF images showed that ERES was indeed greatly recruited to the cis-/medial Golgi and formed an extensive ERES-Golgi interface (Fig. 2E). The association was so evident that most ERES were tightly associated with the cis-Golgi, resulting in much fewer ERES in the periphery of cells (Fig. 2F).

Next, we investigated whether the recruitment between VPS13B and Sec23IP can occur at the endogenous level. Indeed, we found that endogenous Sec23IP was recruited to the Golgi positive for VPS13B-GFP (Fig. 2G, H), as indicated by a significant increase in the percentage of ERES (anti-Sec31A) around the cis-Golgi (anti-GM130), compared to cells without plasmid transfection in the same field of view (Fig. 2G; middle vs bottom panel). On the other hand, the expression of Halo-Sec23IP also increased the association between ERES (anti-Sec31A) and the Golgi (anti-GM130) (Fig. S3E, F).

In addition, a significant proportion of endogenous Sec23IP puncta (∼50%) were associated with endogenous VPS13B at ERES-Golgi interfaces but not at the cell periphery (Fig. 2I, J). This suggested that the interaction between VPS13B and Sec23IP specifically occurred at ERES-Golgi interfaces.

To obtain ultrastructural details about the extensive ERES-Golgi interface mediated by co-expression of VPS13B and Sec23IP, we performed correlative light and electron microscopy (CLEM). We observed that the Golgi appeared to be compacted and fragmented upon the co-expression, and the membranes positive for Halo-Sec23IP appeared to be in the form of small vesicle clusters that were adjacent to the Golgi marked by VPS13B-GFP (Fig. 2K). The cluster of small vesicles in cells overexpressing VPS13B has been observed in our previous study^28^, and this result suggested that these clustered vesicles could be marked by Sec23IP and were possibly derived from ERES. We also observed that several lipid droplets (dark-staining structures; yellow arrows) were present near the Golgi in the cell (Fig. 2K), consistent with our previous study^28^.

Next, we asked whether VPS13B or Sec23IP was required for the formation or maintenance of ERES-Golgi interfaces. The association between the cis-Golgi and ERES was not completely abolished, but resulted in a significant reduction in two independent VPS13B KO clones (Fig. 2L), as shown by the colocalization analyzes (Fig. 2M). In addition, colocalization analysis showed that siRNA-mediated suppression of Sec23IP in VPS13B KO cells reduced the ERES-Golgi associations to a higher extent compared to the VPS13B KO (Figs. 2L, M; S3G). These results suggested that VPS13B was necessary for the association between ERES and the Golgi, and other factors besides VPS13B might be also involved.

### VPS13B bound to the NT of Sec23IP via the VAB domain

Next, we investigated the mechanism by which VPS13B interacted with Sec23IP. The VAB domain of VPS13B was diffused in the cytosol (Fig. 3A). However, a significant portion of the VPS13B-VAB was recruited to ERES by Halo-Sec23IP (Fig. 3B). Accordingly, GFP-trap assays confirmed the interaction of GFP-VPS13B-VAB with Halo-Sec23IP (Fig. 3C). In addition, we found that neither the VPS13B-NT nor the PH-like domain were recruited by Halo-Sec23IP (Fig. 3D). These results suggested that VPS13B interacts with Sec23IP via the VAB domain.

**Fig. 3.**
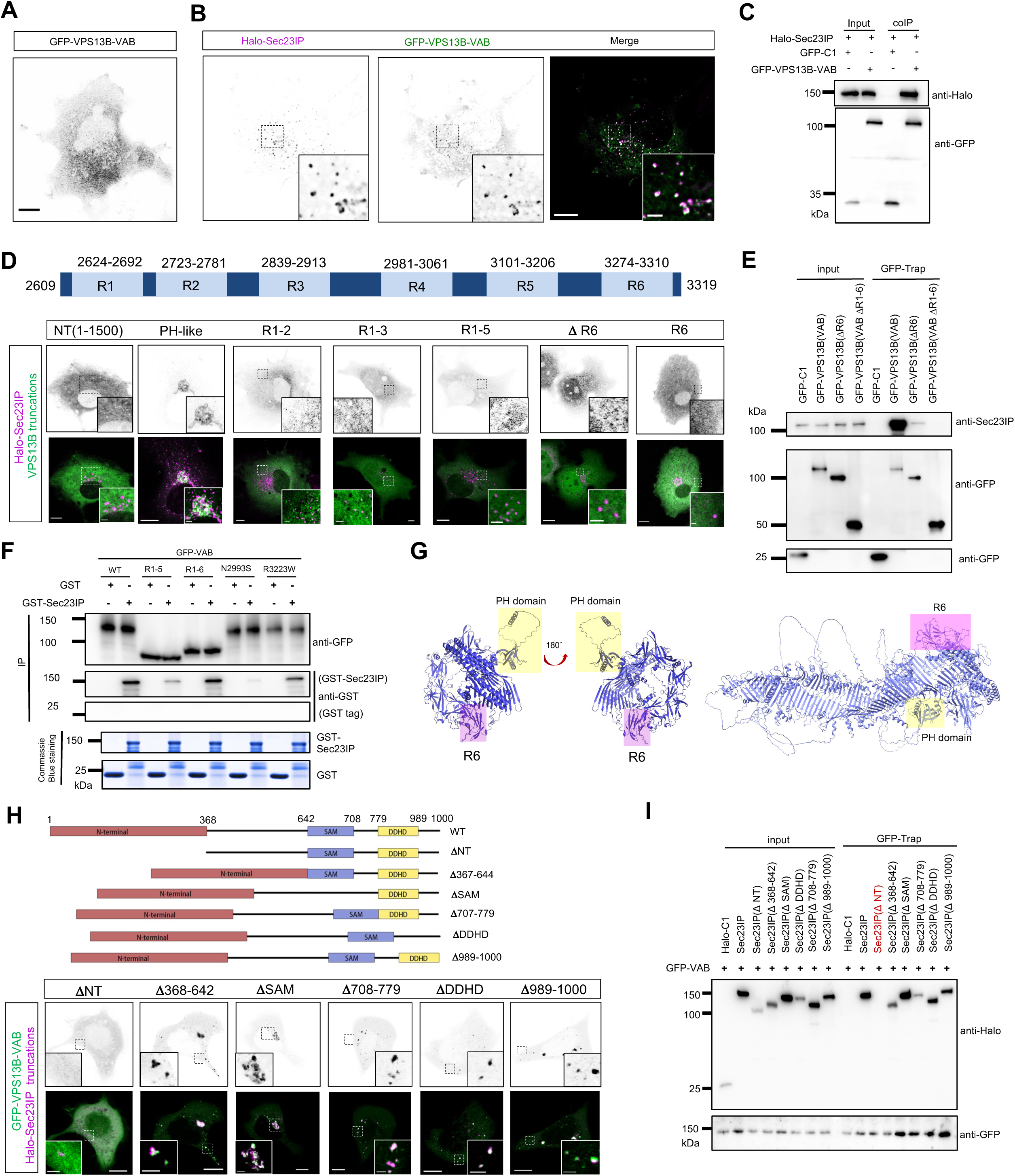
The VAB domain of VPS13B is responsible for the interaction with Sec23IP. (A, B) Representative images of live HeLa cells expressing GFP-VPS13B-VAB (green) alone (**A**) or coexpression GFP-VPS13B-VAB (green) and Halo-Sec23IP (magenta) (**B**) with insets. (C) coIP assays showed an interaction between GFP-VPS13B-VAB and Halo-Sec23IP in HEK293 cells. Blue arrows denoted VPS13B while red arrows indicated Sec23IP in the blots. (D) Top: schematic diagram of six repeats in the VAB domain of VPS13B. Bottom: representative images of live HeLa cells expressing GFP-VPS13B truncations (green) and Halo-Sec23IP (magenta) with insets. (E) coIP assays showed interactions between GFP-VPS13B-VAB truncations and Halo-Sec23IP in HEK293 cells. (F) Pulldown assays showed a direct interaction between GFP-VPS13B-VAB, GFP-R1-6, GFP-R1-5 or two disease-associated missense mutants (N2993S and R3223W) and purified GST-Sec23IP. (G) Alphafold-predicted structures of VPS13B-CT (left panel) or full-length VPS13B (right panel) with the PH domain (yellow) and the VAB (blue). The R6 was highlighted in red. (H) Top: domain organization of Sec23IP. Bottom: representative images of live HeLa cells expressing Halo-Sec23IP truncations (magenta) and GFP-VPS13B-VAB (green) with insets. (I) coIP assays showed interactions between Halo-Sec23IP truncations and GFP-VPS13B-VAB in HEK293 cells. Scale bar, 10 μm in the whole cell images and 2 μm in the insets in A, B, D and H.

The VAB domains of mammalian VPS13 proteins contained six repeats (R1-R6) (Fig. 3D, top panel)^30^. By dissecting the VPS13B-VAB, we found that the sixth repeat (R6) was important for the recruitment because the recruitment was abolished in VPS13B-VAB truncations without R6 (Fig. 3D, bottom panel). However, the R6 alone was unable to be recruited by Sec23IP (Fig. 3D, bottom panel), suggesting that the R6 of the VPS13B-VAB domain is required but not sufficient for the recruitment by Sec23IP. In accord with the imaging results, GFP-trap assays confirmed the crucial role of R6 in the interaction between GFP-VPS13B-VAB and endogenous Sec23IP (Fig. 3E).

We further asked whether the VPS13B-VAB directly interacts with Sec23IP by performing *in vitro* pulldown assays. In this assay, we used GFP-trap to pellet GFP-VPS13B-VAB from HEK293 cells transiently expressing GFP-VPS13B-VAB by using a high-salt (500 mM NaCl) lysis buffer, as described previously^36^. After rigorous washing to remove proteins that could copellet with GFP-VPS13B-VAB under high-salt conditions, GFP-VPS13B-VAB beads were incubated with purified glutathione S-transferase (GST) tag alone or with GST-Sec23IP, respectively. Indeed, GFP-VPS13B-VAB bound to GST-Sec23IP but not to the GST tag (Fig. 3F), indicating that the VAB domain of VPS13B bound to Sec23IP. Consistently, the in vitro pulldown assays also demonstrated that the binding of a region containing R1-5 to His-Sec23IP was significantly weaker than either a region containing R1-6 or the VPS13B-VAB (Fig. 3F), confirming the important role of R6 in the binding. Collectively, these results suggested that the R6 of VPS13B-VAB mediated the interaction with Sec23IP on ERES, meanwhile the PH domain recognized the cis-/medial Golgi via binding toPI4P (Fig. 3G).

Next, we asked how Sec23IP interacted with VPS13B. Sec23IP harbored an NT region, a sterile alpha motif (SAM), and a DDHD domain (Fig. 3H, top panel). We found that deletion of the NT region, but not other domains, abolished the colocalization with the VPS13B-VAB (Fig. 3H, bottom panel), suggesting that the NT of Sec23IP was necessary for the interaction with VPS13B-VAB. This result was further confirmed by GFP-trap assays (Fig. 3I). Considering that Sec23IP also interacted with Sec31A via its NT^35^, we tested whether VPS13B and Sec31A competed with each other for binding to Sec23IP at ERES. Sec23IP was able to colocalize with VPS13B-VAB and Sec31A (Fig. S3H, I), suggesting that Sec23IP could simultaneously bind to VPS13B and Sec31A at ERES.

### Cohen syndrome-associated missense mutations in the VPS13B-VAB impaired the interaction with Sec23IP

Importantly, several missense mutants from Cohen syndrome patients were found in the VAB domain of VPS13B^37^. We therefore systematically examined whether these pathogenic mutations in the VAB domain had impacts on the interaction between VPS13B and Sec23IP (Fig. 4A). Our colocalization analyzes showed that most of these VAB mutants interacted with Sec23IP to a lesser extent than the WT VAB (Fig. 4A-M). Among these mutants, G2729R and N2993S, most significantly impaired the interaction with Sec23IP, as shown by the colocalization analyzes (Fig. 4N).

**Fig. 4.**
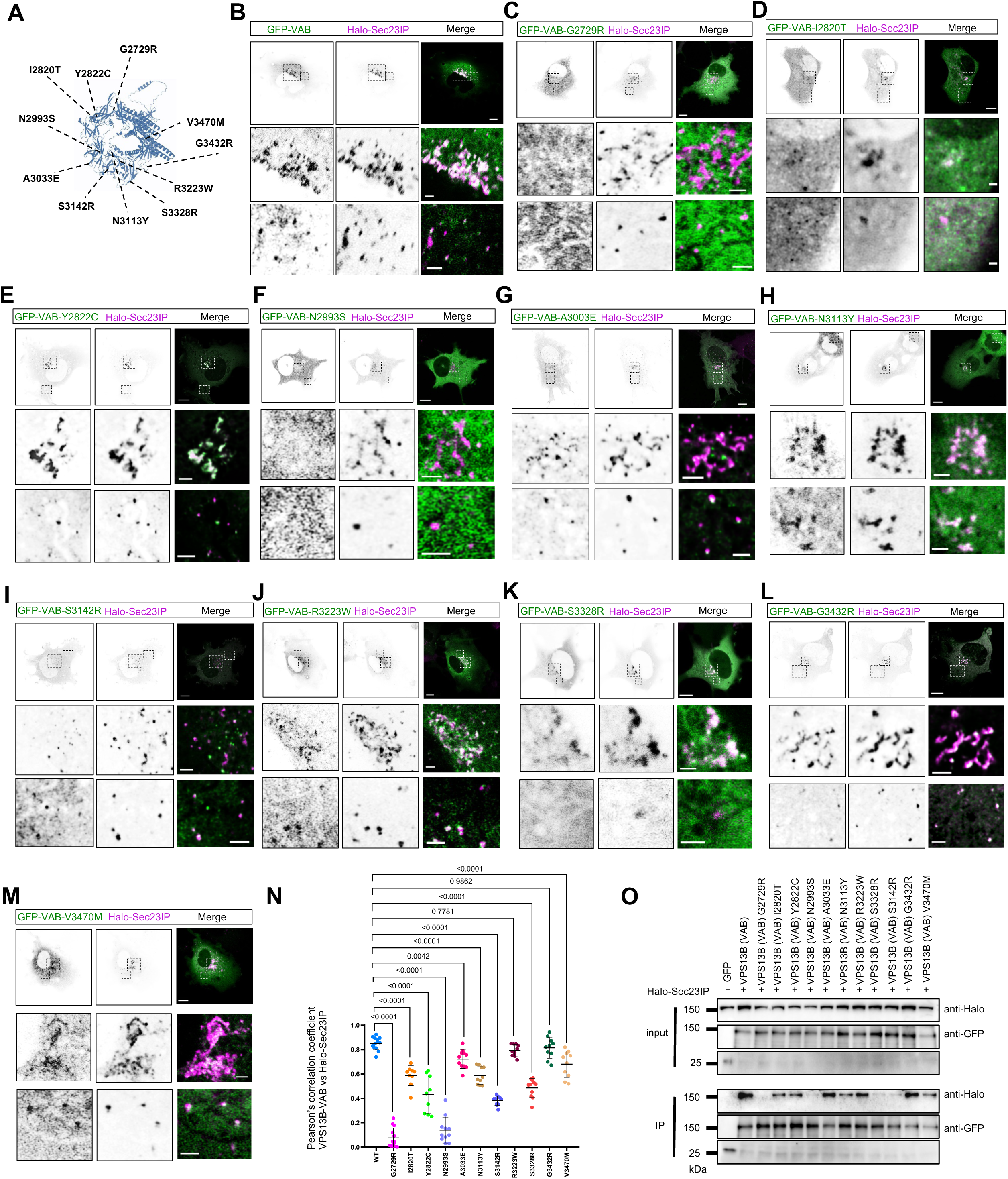
Cohen disease-associated point mutants on VPS13B-VAB domain impaired the interaction with Sec23IP. (A) The disease-associated missense mutants indicated in the Alphafold-predicted structures of VPS13B-CT. (B-M) Representative images of live HeLa cells expressing GFP-VPS13B-VAB containing these disease-associated mutants (green) and Halo-Sec23IP (magenta) with two insets on the bottom. (N) Pearson’s correlation coefficient of disease mutants of GFP-VPS13B-VAB vs Halo-Sec23IP as shown in (B-M) in 3 independent experiments. Ordinary one-way ANOVA with Tukey’s multiple comparisons test. Mean ± SD. (O) coIP assays showed interactions between these disease mutants of GFP-VPS13B-VAB and Halo-Sec23IP as in (B-M) in HEK293 cells. Scale bar, 10 μm in the whole cell images and 2 μm in the insets in B-M.

Accordingly, GFP-trap assays demonstrated that these pathogenic point mutations inhibited the interaction with Sec23IP (Fig. 4O). Among these mutants, G2729R, N2993S, S3142R and S3328R almost completely inhibited the interaction with Sec23IP (Fig. 4O). In addition, in vitro pulldown assays showed that N2993S greatly reduced the binding with purified His-Sec23IP while the other mutant R3323W only had a moderate effect (Fig. 3F). Importantly, most pathogneic mutations of VPS13B were nonsense mutants that resulted in premature termination and the absence of the VAB domain^12, 37^, eventually leading to the loss of the ability to interact with Sec23IP. All together, these results suggested a link between the VPS13B-Sec23IP interaction and Cohen disease, and a defect in the interaction may contribute to the pathogenesis of this disease.

### VPS13B was required for the biogenesis of tubular ERGIC

Bridge-like RGB repeating lipid transporters, including Vps13 proteins and Atg2, were thought to play a key role in de novo biogenesis of organelles by providing membrane lipids to meet requirements during membrane expansion and growth^38–43^. Since our results showed that VPS13B interacted with Sec23IP at ERES-Golgi interfaces, we wondered whether VPS13B and Sec23IP were required for the biogenesis of membrane structures in the early secretory pathway. The formation of ERES, marked by either endogenous Sec31A (outer coat; Fig. S4A) or Sec23IP (Fig. S4B) in IF, appeared not to be greatly affected upon VPS13B KO. COPI vesicles labeled by the coatomer subunits COPA was not significantly affected as well (Fig. S4C). Furthermore, the number or size of the conventional tubulo-vesicular ER-Golgi Intermediate Compartment (ERGIC) marked by anti-ERGIC53 appeared not to be strongly altered in VPS13B KO (Fig. S4D).

Importantly, we found that VPS13B KO dramatically impaired the formation of an unconventional tubular ERGIC (tERGIC) (Fig. 5A, B), which specifically accelerated ER-to-Golgi trafficking of certain soluble cargoes^44^. In control cells, expression of GFP-Rab1B induced the formation of tERIGC, which was highly elongated (Fig. 5A). In contrast, VPS13B KO greatly impaired the formation of Rab1B-induced tERGIC (Fig. 5B), resulting in a strong reduction in both the length (Fig.5D) and number (Fig.5E) of tERGIC compared to control. In addition, we found that the formation of tERGIC was also significantly impaired in two independent Sec23IP KO clones (Fig. 5C-E). Together, these results indicated that VPS13B and its binding partner Sec23IP were indispensable for tERGIC formation.

**Fig. 5.**
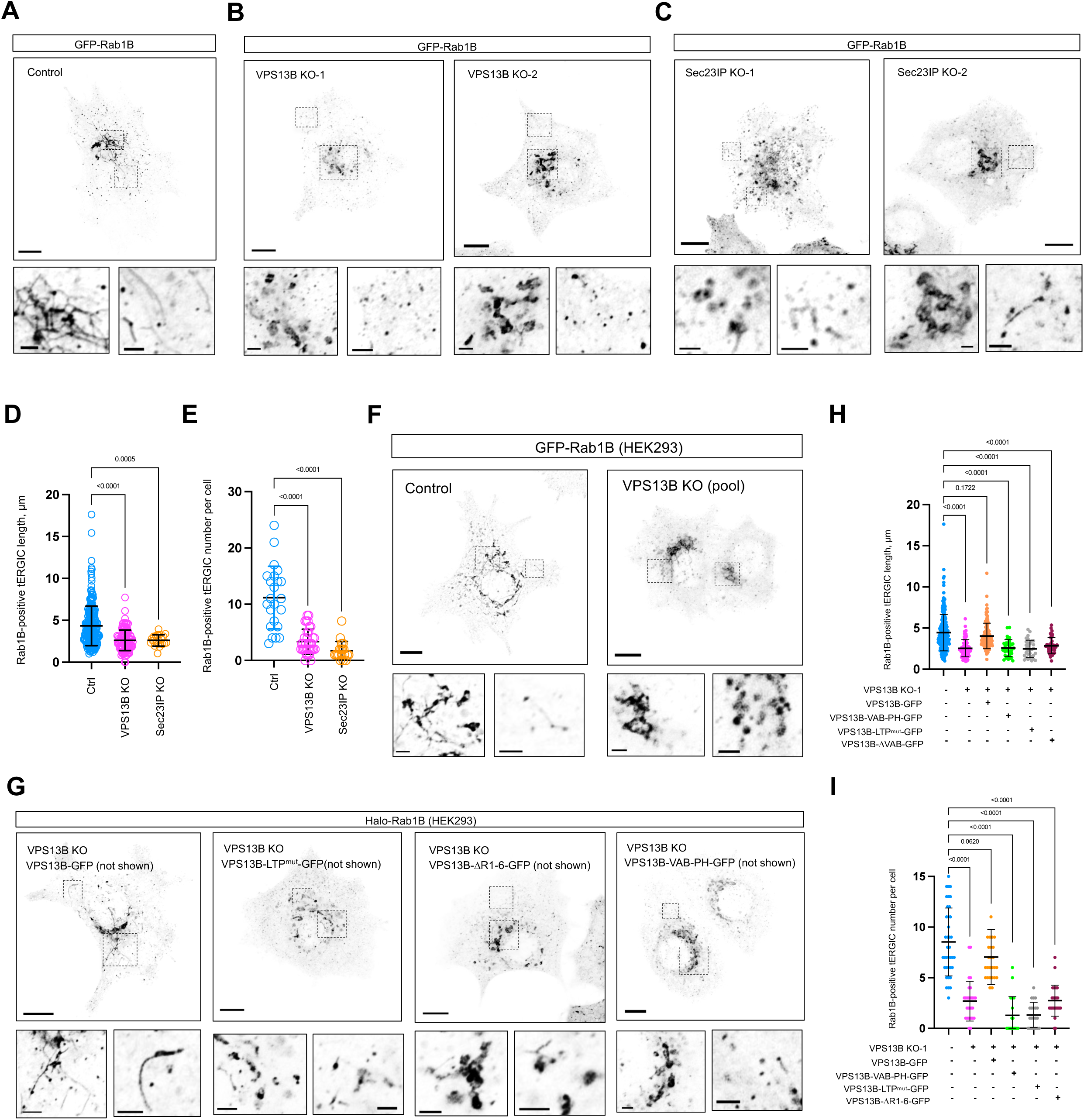
VPS13B KO blocks the formation of tubular ERGIC. (A-C) Representative images of live control HeLa (**A**), two VPS13B KO clones (**B**) or two Sec23IP KO clones expressing GFP-Rab1B with two insets on the bottom. The left inset was from the Golgi region and the right inset was from the cell periphery. (D, E) The length (**D**) or number (**E**) of tERGIC labeled by GFP-Rab1B. N=257 from 23 control cells, N= 120 from 35 VPS13B KO cells and N= 20 from 19 Sec23IP KO cells in 3 independent experiments. Mean ± SD. Two tailed unpaired student t test. (F) Representative images of a VPS13B-KO HEK293 pool expressing GFP-Rab1B with two insets on the bottom. (G) Representative images of a VPS13B-KO HEK293 pool expressing either WT VPS13B-GFP, VPS13B-ΔVAB-GFP, VPS13B-VAB-PH-GFP and a potential lipid transfer deficient mutant (VPS13B-LTP^mut^-GFP) along with Halo-Rab1B with two insets on the bottom. (H, I) The length (H) and number (I) of GFP-Rab1B tERGIC in (F, G). More than 20 cells were quantified for each condition from 3 independent experiments. Ordinary one-way ANOVA with Tukey’s multiple comparisons test. Mean ± SD. Scale bar, 10 μm in the whole cell images and 2 μm in the insets in A, B, C, F and G.

Next, we investigated the mechanisms underlying the role of VPS13B in tERGIC formation by performing rescue experiments. The transfection efficiency of the VPS13B construct and its mutants in VPS13B KO HeLa cells was low and unsuitable for the rescue experiments. Therefore, we generated a pool of VPS13B-KO HEK293 cells using CRISPR-Cas9 (Fig. S5A, B). The tERGIC defect was robustly observed in the VPS13B-KO HEK293 cell pool (Fig. 5F). Remarkably, introduction of WT VPS13B-GFP significantly rescued the tERGIC phenotype (Fig. 5G-I), indicating that the defect in the tERGIC formation was specific to VPS13B.

Next, we investigated whether and to what extent the formation of tERGIC was dependent on the lipid transfer activity of VPS13B. First, we made a lipid transfer-deficient mutant (VPS13B-LTP^mut^-GFP), in which a few hydrophobic residues in the midway of the hydrophobic groove of VPS13B were mutated to hydrophilic residues to block lipid transport according to a recent study on Vps13 (Fig. S5C)^45^. Importantly, the lipid-transfer-deficient mutant was unable to rescue the defect in tERGIC formation resulted from VPS13B KO (Fig. 5F-H), suggesting that lipid transfer of VPS13B was indispensible for tERGIC formation.

In addition, we found that the introduction of a Sec23IP binding defective VPS13B mutant (VPS13B-ΔR1-6) into VPS13B KO HEK293 cells was unable to fully rescue the phenotype (Fig. 5F-H). Interestingly, a truncated mutant only containing the VAB and PH domain was also unable to restore the defect (Fig. 5F-H). Together, our findings suggested that both the VAB domain and the lipid transfer activity were indispensable for the process.

### VPS13B and Sec23IP promoted the ER export of procollagen

One of cardinal features in patients with Cohen syndrome was joint hypermobility, which was linked to defects in collagen biogenesis and/or secretion^46^. Therefore, we asked whether VPS13B played a role in ER-Golgi trafficking of procollagen, the most abundant protein in human body. We tracked the ER-to-Golgi trafficking of procollagen using primary mouse embryonic fibroblasts (MEF) as a cell model in IF. In the assays, we used an antibody (SP1.D8) that specifically recognized intracellular procollagen IA. The release of procollagen from the ER was synchronized by the addition of ascorbic acid, which was a key factor for hydroxyproline formation in procollagen^47^. As a control, ER exit of procollagen IA was substantially delayed by treatment with bredfeldin A (BFA), an ER-Golgi trafficking inhibitor (Fig. 6D). Importantly, depletion of VPS13B caused accumulation of procollagen in the ER prior to release (Fig. 6E) and resulted in a significant delay in ER export of procollagen compared to the control (Fig. 6A, B), as shown by the changes in intracellular procollagen IA fluorescence over time (Fig. 6F). Surprisingly, depletion of Sec23IP inhibited ER export of procollagen to an even higher extent than VPS13B depletion (Fig. 6C, F & G), indicating that Sec23IP acted as an important factor for procollagen secretion, and Sec23IP may regulate ER export of procollagen via other pathways besides the recruitment of VPS13B, for instance, by modulation of the organization of ERES^34, 35^.

**Fig. 6.**
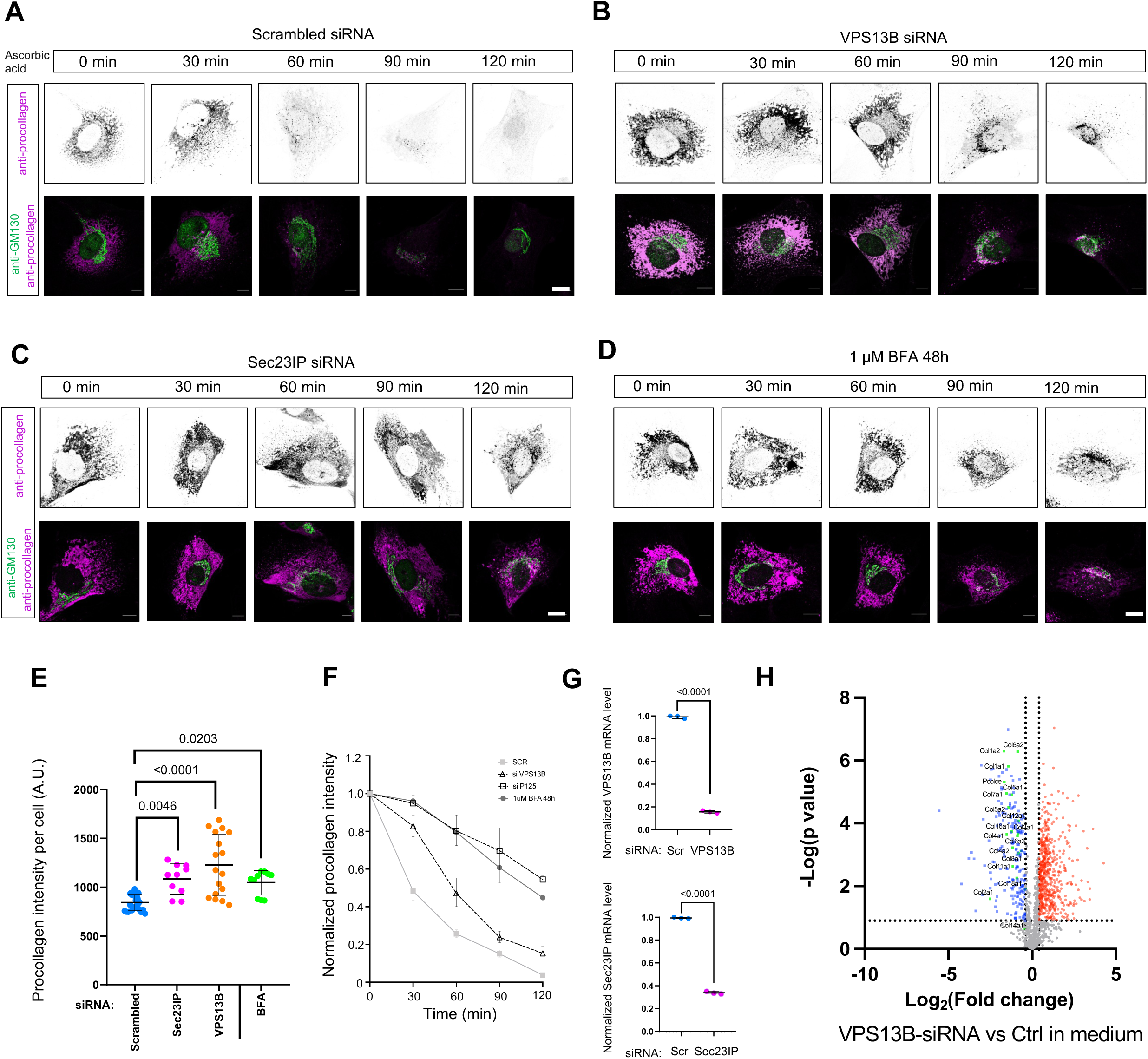
Depletion of VPS13B or Sec23IP reduces ER exit of procollagen. (A-D) Representative confocal images of fixed control (A), VPS13 depleted (B), Sec23IP depleted (C) primary MEF cells labeled with anti-procollagen (magenta) and anti-GM130 (green) upon ascorbic acid (50 μM) stimulation with five timepoints (0, 30 min, 60 min, 90 min, 120 min). In this assay, control primary MEF cells pre-treated with BFA (1 μM; 48 h) was used as a control (D). (E) The fluorescence intensity of anti-procollagen prior to ascorbic acid stimulation. 23 control cells and 17 from xxx VPS13B KO cells in 3 independent experiments. Mean ± SD. Ordinary one-way ANOVA with Tukey’s multiple comparisons test. (F) The changes of fluorescence intensity of anti-procollagen upon ascorbic acid stimulation as in (A-D). 18 control cells and 19 VPS13B KO cells in 3 independent experiments. Mean ± SD. Ordinary one-way ANOVA with Tukey’s multiple comparisons test. (G) qPCR assays indicated the efficiency of shRNA-mediated suppression of VPS13B (top panel) and Sec23IP (bottom panel) in primary MEF cells from 3 independent experiments. Mean ± SD. Two tailed unpaired student t test. (H) Volcano plot of proteins in the medium of control compared to VPS13B-depleted primary MEF cells. Candidates that were considered significant (−log [P value] > 1.3; P < 0.05) were labeled in orange (Log2 [fold change] > 0; increased in abundance) or blue (Log2 [fold change] < 0; decreased in abundance). All collagens identified in the quantitative MS were labeled. Scale bar, 10 μm in (A-D).

To further confirm the role of VPS13B in collagen secretion, we examined the secreted collagens in the medium of primary MEFs synchronzied by ascorbic acid by quantitative mass spectrometry. Several collagen species, including collagen IA, were successfully identified in our MS assays (Fig. 6H). Importantly, the levels of all these collagens were signifcaintly reduced by VPS13B depletion compared to the control group (Fig. 6H). Of note, we found that VPS13B depletion led to aberrent increases in secretion of some proteins, which may be due to indirect consequences of disorganized tERGIC or the Golgi resulting from VPS13B depletion. Taken together, these results suggested that VPS13B and Sec23IP were novel regulatory factors for ER export of procollagens.

## Discussion

Our study reveals a crucial interaction between VPS13B and Sec23IP that promotes the formation of ERES-Golgi interfaces. Cohen syndrome-associated missense mutations in the VAB domain of VPS13B impair the interaction with Sec23IP. VPS13B KO abolishes the formation of tERGIC, the unconventional ER-to-Golgi cargo carrier. Using primary MEF cells as a system to study collagen secretion, we found that depletion of VPS13B significantly delays ER export of procollagens, linking procollagen secretion to joint laxity in patients with Cohen disease. Taken together, we hypothesized that loss of VPS13B-Sec23IP interactions blocked the biogenesis of tERGIC membranes, reducing the efficiency of membrane trafficking and secretion and/or trafficking of proteins/lipids required for the nervous system and joint development at certain developmental stages, ultimately contributing to the pathogenesis of Cohen syndrome (Fig. S5D; working model).

VPS13B was reported to play an essential role in the formation of acrosomes during sperm development in mice^27^, but the molecular mechanisms were unclear. Interestingly, Sec23IP, the VPS13B adaptor identified in this study, was also required for acrosome biogenesis^33^. Therefore, we speculated that the VPS13B-Sec23IP interaction at ERES-Golgi interface may play an important role in acrosome formation in spermiogenesis. Overall, our findings revealed a conserved mechanism in early secretory pathway for Cohen disease pathogenesis and acrosome biogenesis, mechanistically linking the two previously unrelated cellular processes. Given that collagen was not required for acrosome biogenesis, we speculated that VPS13B and its interactor Sec23IP might expedite the ER-to-Golgi trafficking of a wide range of proteins and lipids, but not a certain type of cargo, in response to developmental signals.

While the cellular functions of the early secretory pathway were generally conserved among all eukaryotes, the organization of the ER–Golgi interface varied widely among species^1^. For example, plants and some yeast species had a compact organization of the ER and Golgi, in which Golgi mini stacks were dispersed throughout the cytosol and moved along actin cables adjacent to the ER. In contrast to the scattered ministacks, the Golgi of animal cells was present in the form of a ribbon that assemble in the perinuclear. Interestingly, deletion of VPS13B or Sec23IP in mammalian cells resulted in dispersed Golgi mini stacks over the cytosol, which is similar to the organization of ER and Golgi in plants and yeasts. Since both VPS13B and Sec23IP had no ortholog in plants and yeasts, VPS13B and Sec23IP may be responsible for the unique ER-Golgi organization in mammals.

It is controversial how procollagen was exported from the ER. Newly synthesized procollagen was exported from the ERES under the control of a small GTPase Sar1 in coordination with other factors, including TANGO1^48, 49^, cTAGE5^50^, Seldin1^51^, KLHL12^52^. Due to the size of procollagen triple helices, it was assumed that they were transported from the ER to the Golgi in specialized large COPII-dependent carriers (>300 nm). However, in animal cells, the traditional vesicle carrier model was challenged by the recent nano-resolved structure of ERES identified as a continuous network of interwoven membrane tubules connecting the ER and extruding pearl-shaped extensions towards the Golgi^53^. Furthermore, in a recent study, McCaughey et al. showed that transport of procollagen to the Golgi may not involve long-range transport of large vesicular structures, and thus proposed a short-loop model of COPII-dependent transport that facilitated local transfer of procollagen from the juxtanuclear ER to the Golgi through the local formation of budding structures at ERES in close proximity to Golgi membranes^54^. In this study, our results show that the VPS13B-Sec23IP interaction promotes ERES-Golgi interfaces and facilitates ER export of procollagens, supporting the notion that ER export of procollagen may not require large vesicles. Furthermore, our findings were also in line with models of ERGIC-dependent expansion of COPII carriers^55, 56^, in which procollagen did not utilize vesicles during transport between Golgi stacks but remained within cisternae^57^.

Notewhorthy, depletion of Sec23IP or VPS13B did not completely block ER export of procollagen. One possibility was that the efficiency of siRNA-mediated knockdown is not 100%. However, we argue that it is probably not the case because both VPS13B and Sec23IP were absent in *Caenorhabditis elegans*, but collagen secretion was intact in this organism (unpublished data). Clinically, although Cohen dyndrome was characterized by developmental delays and joint hypermotility, patients suffered from this disease could survive and grow up, indicating that the syntheis and secretion of collagen may be impacted but not turned off when the VPS13B or Sec23IP gene was mutated. We therefore speculated that VPS13B was not necessarily required for basal ER export of procollagen, but it could accelerate its secretion under certain functional contexts, for instance, at certain development stages or in certain tissues that required high intensity of procollagen secretion. Supporting the notion, VPS13B loss of function was not lethal in mouse or in cell cultures, but was required for neurite outgrowth and acrosome biogenesis, both of which required the high intensity of protein and lipid trafficking. In addition, tERGIC was not always present, but its biogenesis was induced either by Rab1 activity or overload of certain types of cargoes^44^, both stimuli could be considered as cellular stresses impacted on the early secretory pathway. This suggested that VPS13B may also function in a stress-induced manner. Another possible explanation was that cells can compensate for the loss of VPS13B by upregulating other proteins. For instance, VPS13D, the paralog of VPS13B in mammals, may play a moonlighting role in the early secretory pathway, as previous studies showed that part of VPS13D was localized in the Golgi^21,41^. Further studies are needed to investigate the functional relationship between VPS13 proteins in procollagen trafficking.

Another question not answered in this study was whether tERGIC carried procollagen triple helices. tERGIC was a highly elongated tubular membrane, with a length of 2–20 μm and a diameter of <30 nm^44^. Meanwhile, a procollagen triple helix was ∼300 nm long and ∼1.5 nm in diameter. Theoretically, tERGIC was an optimal carrier for procollagen triple helices because of its high surface-to-volume ratio, high intracellular movement speed, and ER-Golgi recycling abilities^44^. However, we could not observe the evident appearance of tERGIC during the tracking of procollagen secretion in primary MEFs in IF because the fixation process would disrupt the elongated thin membranes of tERGIC (unpublished results), which made the observation of tERGIC technically difficult. Nevertheless, we cannot exclude the possibility that tERGIC indirectly regulated procollagen trafficking because the high ER-Golgi recycling capabilities of tERGIC may regulate the protein or lipid compositions of the COPII complex specialized for procollagen secretion. Overall, our study clearly demonstrated specific and important roles of VPS13B and Sec23IP in ER export of procollagen and establishes a link between procollagen secretion and joint laxity in patients suffered from Cohen disease.

## METHODS

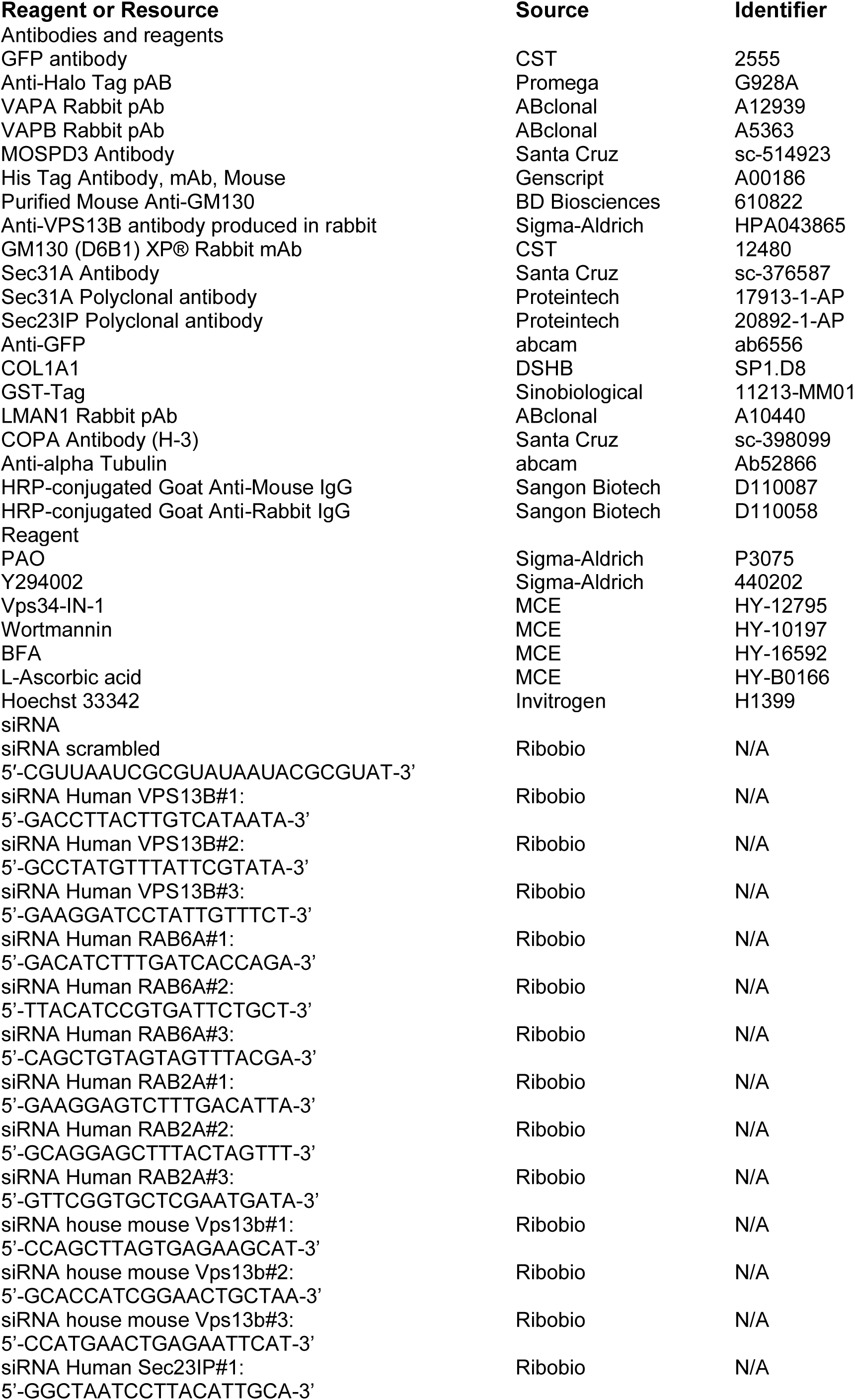
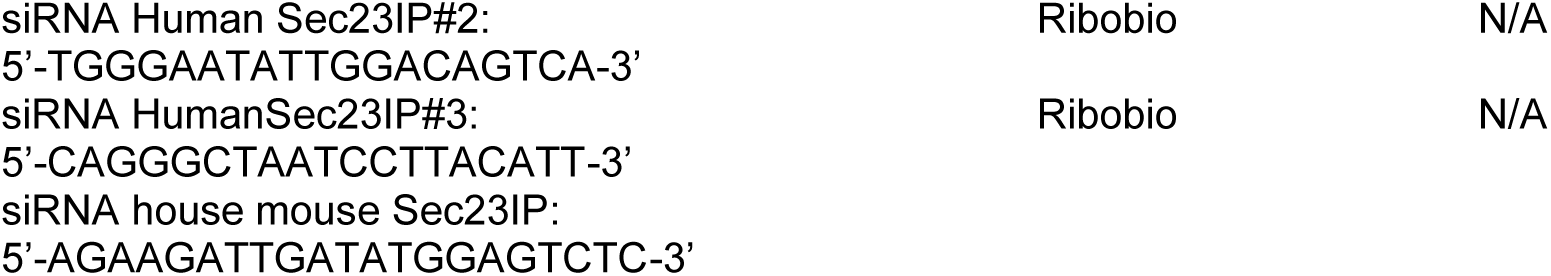

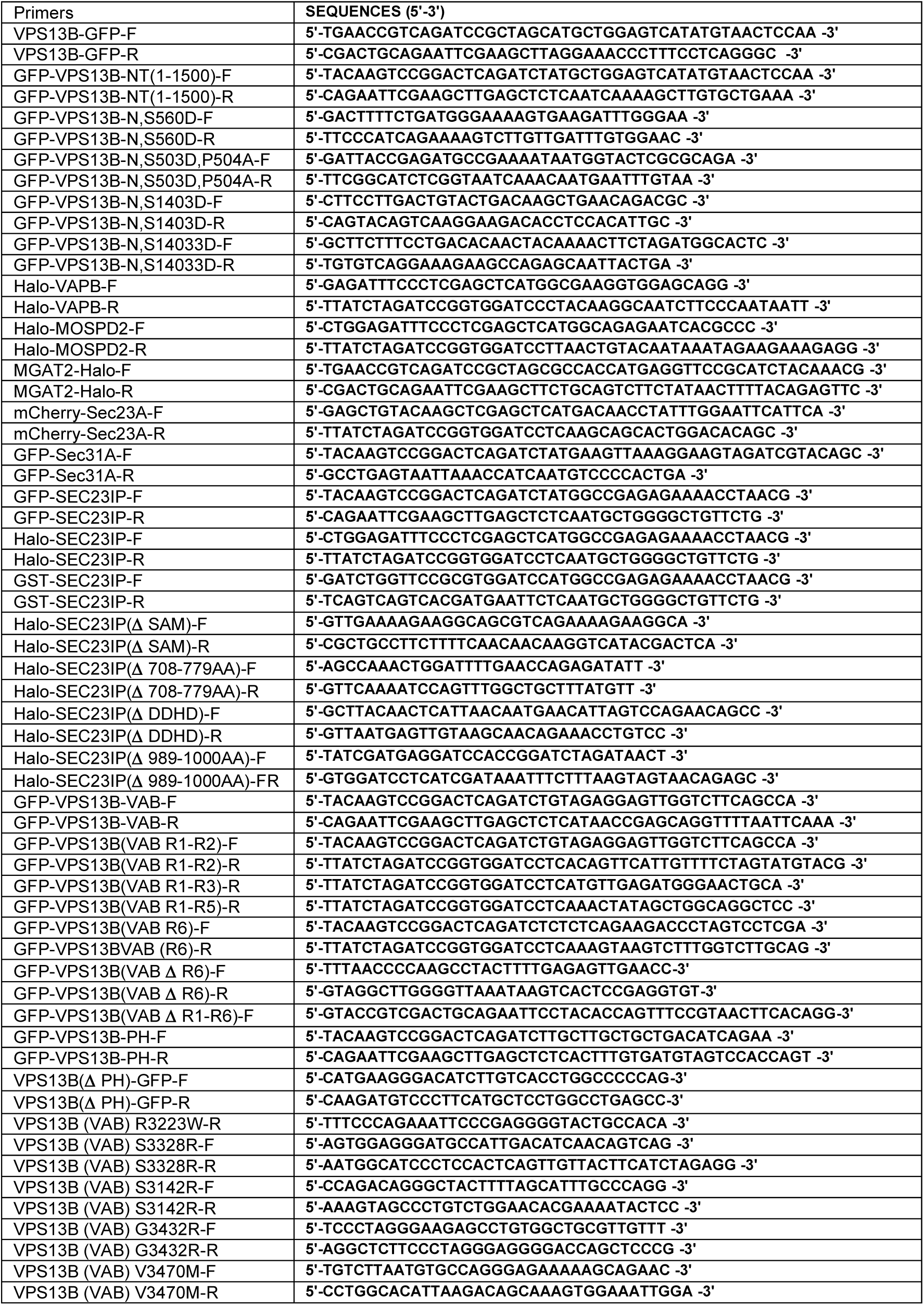
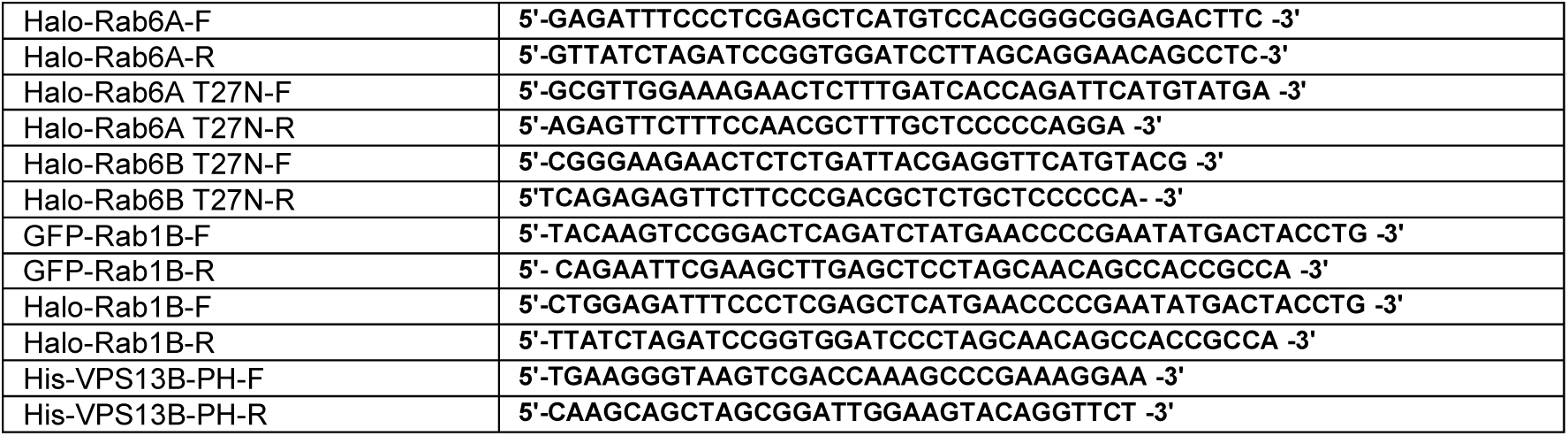

### Cell culture, transfection, RNAi

The African green monkey kidney fibroblast-like COS7 cell line (ATCC), human cervical cancer HeLa cells (ATCC), and human embryonic kidney 293T (ATCC) were grown in (Invitrogen) supplemented with 10% fetal bovine serum (Gibco). All of the cell lines used in this study were confirmed free of mycoplasma contamination.

Mouse embryos were removed from pregnant C57BL/6 mice at day E13.5. The head, limb buds, and visible internal organs were removed from the embryo, followed by the treatment of 500 μL trypsin (0.05%) for 30 min at 37°C. The trypsin was inactivated using an equal volume of DMEM supplemented with 10% FBS, and the mixture was pipetted up and down several times to dissociate the embryo into single cells. The cells were suspended in DMEM supplemented with 10% FBS and 1% penicillin/streptomycin, and then were seeded into 10-cm culture dishes. The dishes were incubated under standard cell culture conditions at 37°C, 5% CO2.

Transfection of plasmids and RNAi oligos was carried out with Lipofectamine 2000 and RNAi MAX, respectively. For transfection, cells were seeded at 4 x 10^5^ cells per well in a six-well dish ∼16 h before transfection. Plasmid transfections were performed in OPTI-MEM (Invitrogen) with 2 μl Lipofectamine 2000 per well for 6 h, followed by trypsinization and replating onto glass-bottom confocal dishes at ∼3.5 x 10^5^ cells per well. Cells were imaged in live-cell medium (DMEM with 10% FBS and 20 mM Hepes with no penicillin or streptomycin) ∼16–24 h after transfection. For siRNA transfections, cells were plated on 3.5 cm dishes at 30–40% density, and 2 μl Lipofectamine RNAimax (Invitrogen) and 50 ng siRNA were used per well. At 48 h after transfection, a second round of transfection was performed with 50 ng siRNAs. Cells were analyzed 24 h after the second transfection for suppression.

### CRISPR-Cas9-mediated gene editing

To make VPS13B KO HeLa or HEK293 cell lines, two gRNAs (5-AGACGTGACAGCTAGAGTGG-3 and 5-CTAGTGACTCTAGGTCAACA-3) were used to delete ∼76 bp from exon23 of VPS13B gene. To make Sec23IP KO HeLa, two gRNAs (5-TATGGATTGTATCCTGGTTG-3 and 5-ATTAGCCCTGCTGCTGCCAG-3) were used to delete ∼31bp from exon 2 of Sec23IP gene. Complementary gRNAs were annealed and subcloned into the pSpCas9(BB)-2A-GFP (pX-458) vector (#48138; Addgene) between BbsI endonuclease restriction sites. Upon transfection, cells were grown in an antibiotic-free medium for 48 h, followed by single-cell sorting by fluorescence-based flow cytometry.

### GFP-trap assay

GFP trap (GTA-100; ChromoTek) was used for the detection of protein–protein interactions, and the GFP-Trap assays were performed according to the manufacturer’s protocol. Briefly, after 24 h transfection with the indicated plasmids, cells were lysed in ice-cold lysis buffer (50 mM Tris-HCl, pH 7.5, 150 mM NaCl, 1 mM EDTA, 1% Triton X-100 and protease inhibitor cocktail). Lysates were centrifuged at 13,000 rpm for 10 min at 4°C and pellets were removed. Supernatants were incubated with GFP-Trap agarose beads for 1 h at 4°C with gentle shaking. After washing four times with lysis buffer, beads were boiled with SDS sample buffer. Proteins of interest were analyzed by immunoblotting. 5% input was used in GFP traps unless otherwise indicated.

### Protein purification

GST and His constructs were transformed into Escherichia coli BL21 (DE3) cells, and cells were incubated at 37°C until the optical density (OD) at 600 nm reached 0.6–0.8. Subsequently, cells were incubated at 16°C for another hour, followed by induction with 1 mM IPTG overnight at 16°C.Cells were lysed via sonication. GST fusion proteins were purified via the GST-tag Protein Purification kit (C600031-0025, Sangon, China), His fusion proteins were purified via the Ni-NTA Sefinose (TM) Resin Purification kit (G600033-0100, Sangon, China).

### In vitro Pull-down assays of GFP-VPS13B(VAB) and GST-Sec23IP

HEK293 cells transiently transfected with GFP-VPS13B(VAB)were lysed in high-salt lysis buffer (RIPA buffer containing 500 mM NaCl, proteasome inhibitors and PMSF). GFP-Trap beads were used to pellet GFP-VPS13B(VAB) from cell lysates, followed by washing with high-salt lysis buffer for 10 times. The GFP-VPS13B(VAB) beads were incubated with Purified GST–SEC23IP or GST-only overnight at 4°C, respectively, followed by washing beads with freshly prepared HNM buffer (20 mM Hepes, pH 7.4, 0.1 M NaCl, 5 mM MgCl2, 1 mM DTT and 0.2% NP-40). GFP-VPS13B(VAB) beads were resuspended in 100 μL 2 x SDS-sampling buffer. Re-suspended beads were boiled for 10 min at 95°C to dissociate protein complexes from beads. Western blotting was performed using anti-GFP, GST or Sec23IP antibodies. The Coomassie staining was performed for purified GST-Sec23IP.

### PIP Strip assays

The PIP Strips (P-6001) were blocked by TBS-T + 3% fatty acid–free BSA, and then were gently agitated for 1 h at room temperature, followed by an incubation with purified His-VPS13B-PH (0.5 µg/mL) in TBS-T + 3% fatty acid–free BSA overnight at 4°C. After washing the PIP Strips with TBS-T + 3%fatty acid–free BSA three times under gentle agitation for 10 min each time, PIP strips were incubated with the anti-His antibodies overnight at 4°C, followed by repeated washing steps.

### Cryosectioning, immunolabeling and electron microscopy

We followed the Tokuyasu method as described previously^58^. HeLa cells were fixed in 2% formaldehyde and 0.01% glutaraldehyde in PB buffer at 4°C overnight and then washed with pre-cold PB /Glycine. The cells were scraped from the bottom of the plastic dishes in 1% gelatin of PB buffer, centrifuged at 1000rpm/min for 2min and suspended in 12% gelatin at 37°C for 10min. The gelatin-cell mixture was solidified on ice for 15min. Small blocks about 0.5 mm^3^ were made and immersed in 2.3M sucrose overnight at 4“C. Cryosections of 70 nm were made at 120°C with an ultratome (Leica EM FC7). After sections were thawed at room temperature, immunolabeling was performed with anti-VPS13B antibodies followed by immune-Gold secondary antibody. The sections were treated with methyl cellulose/uranyl acetate and subsequently imaged under the H-7650 80kv transmission electron microscope.

### Live imaging by high-resolution confocal microscopy

Cells were grown on 35 mm glass-bottom confocal MatTek dishes, and the dishes were loaded to a laser scanning confocal microscope (LSM980, Zeiss, Germany) equipped with multiple excitation lasers (405 nm, 458 nm, 488 nm, 514 nm, 561 nm and 633 nm) and a spectral fluorescence GaAsP array detector. Cells were imaged with the 63×1.4 NA iPlan-

Apochromat 63 x oil objective using the 405 nm laser for BFP, 488 nm for GFP, 561nm for OFP, tagRFP or mCherry and 633nm for Janilia Fluo® 646 HaloTag® Ligand.

### Mass spectrometry for identification of VPS13B-interacting proteins

The identification of VPS13B-GFP interacting proteins by MS was described in our previous study^59^. Briefly, the bound proteins were extracted from GFP-Trap agarose beads using SDT lysis buffer (4% SDS, 100 mM DTT, 100 mM Tris-HCl pH 8.0), followed by sample boiling for 3 min and further ultrasonicated. Undissolved beads were removed by centrifugation at 16,000 g for 15 min. The supernatant, containing proteins, were collected. Protein digestion was performed with FASP method. Briefly, the detergent, DTT and IAA in UA buffer was added to block-reduced cysteine. Finally, the protein suspension was digested with 2 µg trypsin (Promega) overnight at 37°C. The peptide was collected by centrifugation at 16,000 g for 15 min. The peptide was desalted with C18 StageTip for further LC-MS analysis. LC-MS/MS experiments were performed on a Q Exactive Plus mass spectrometer that was coupled to an Easy nLC (Thermo Fisher Scientific). Peptide was first loaded to a trap column (100 µm x 20 mm, 5 µm, C18, Dr Maisch GmbH, Ammerbuch, Germany) in buffer A (0.1% formic acid in water). Reverse-phase high-performance liquid chromatography (RP-HPLC) separation was performed using a self-packed column (75 µm x 150 mm; 3 µm ReproSil-Pur C18 beads, 120 Å, Dr Maisch GmbH, Ammerbuch, Germany) at a flow rate of 300 nl/min. The RP-HPLC mobile phase A was 0.1% formic acid in water, and B was 0.1% formic acid in 95% acetonitrile. The gradient was set as following: 2%–4% buffer B from 0 min to 2 min, 4% to 30% buffer B from 2 min to 47 min, 30% to 45% buffer B from 47 min to 52 min, 45% to 90% buffer B from 52 min and to 54 min, and 90% buffer B kept until to 60 min. MS data was acquired using a data-dependent top20 method dynamically choosing the most abundant precursor ions from the survey scan (350– 1800 m/z) for HCD fragmentation. A lock mass of 445.120025 Da was used as internal standard for mass calibration. The full MS scans were acquired at a resolution of 70,000 at m/z 200, and 17,500 at m/z 200 for MS/MS scan. The maximum injection time was set to 50 ms for MS and 50 ms for MS/ MS. Normalized collision energy was 27 and the isolation window was set to 1.6 Th. Dynamic exclusion duration was 60 s. The MS data were analyzed using MaxQuant software version 1.6.1.0. MS data were searched against the UniProtKB Rattus norvegicus database (36,080 total entries, downloaded 08/14/2018). Trypsin was selected as the digestion enzyme. A maximum of two missed cleavage sites and the mass tolerance of 4.5 ppm for precursor ions and 20 ppm for fragment ions were defined for database search. Carbamidomethylating of cysteines was defined as a fixed modification, while acetylation of protein N-terminal, oxidation of Methionine was set as variable modifications for database searching. The database search results were filtered and exported with a <1% false discovery rate (FDR) at peptide-spectrum-matched level, and protein level, respectively.

### Halo staining in live cell

Cells were incubated with complete medium with 5nM Janilia Fluo® 646 HaloTag® Ligand for 30 minutes. Cells were washed three times with complete medium to remove extra ligands, followed by incubation for another 30 minutes. Medium was replaced with imaging medium to remove unconjugated Halo ligands that has diffused out of the cells prior to imaging.

### Immunofluorescence staining

Cells were fixed with 4% PFA (paraformaldehyde, Sigma) in PBS for 10 min at room temperature. After washing with PBS three times, cells were permeabilized with 0.1% Triton X-100 in PBS for 15 min on ice. Cells were then washed three times with PBS, blocked with 0.5% BSA in PBS for 1 h, incubated with primary antibodies in diluted blocking buffer overnight, and washed with PBS three times. Secondary antibodies were applied for 1 h at room temperature. After washing with PBS three times, samples were mounted on Vectashield (H-1000; Vector Laboratories).

## Image and statistical analysis

All image analysis and processing were performed using ImageJ (National Institutes of Health). All statistical analyses and p-value determinations were performed in GraphPad Prism6. All the error bars represent mean ± SD. To determine p-values, ordinary one-way ANOVA with Tukey’s multiple comparisons test was performed among multiple groups and a two-tailed unpaired student t-test was performed between two groups.

## ACKNOWLEDGEMENTS

We thank Anbing Shi and Yanling Yan (Huazhong University of Science and Technology) for discussions. We thank Ying Li (Cryo-EM Facility of Tsinghua University, Branch of National Protein Science Facility) for cryosection and immunolabeling. We thank the Mass Spectrometry Core Facility (Mr. Cookson K.C.Chiu), the Bio-imaging Core Facility (Dr. Zhenglong Sun and Ms. Mei Yu), and the Sequencing Core Facility (Dr. MiaoCui) of Shenzhen Bay Laboratory for providing technical supports.

## AUTHOR CONTRIBUTIONS

Y. Du, and W. Ji conceived the project and designed the experiments. Y. Du and X. Fan performed the experiments. Y. Du, X. Fan, C. Song, J. Xiong and W. Ji analyzed and interpreted the data. W. Ji prepared the manuscript with inputs and approval from all authors.

## FUNDING

W. Ji was supported by National Natural Science Foundation of China (92354304; 32371343; 32122025). J. Xiong was supported by National Natural Science Foundation of China (81901166).

## Competing interests

The authors declare no competing interests.

## Data and materials availability

All the data and relevant materials, including reagents and primers, that supports the findings of this study are available from the corresponding author upon reasonable request.

---

**Fig. S1.**
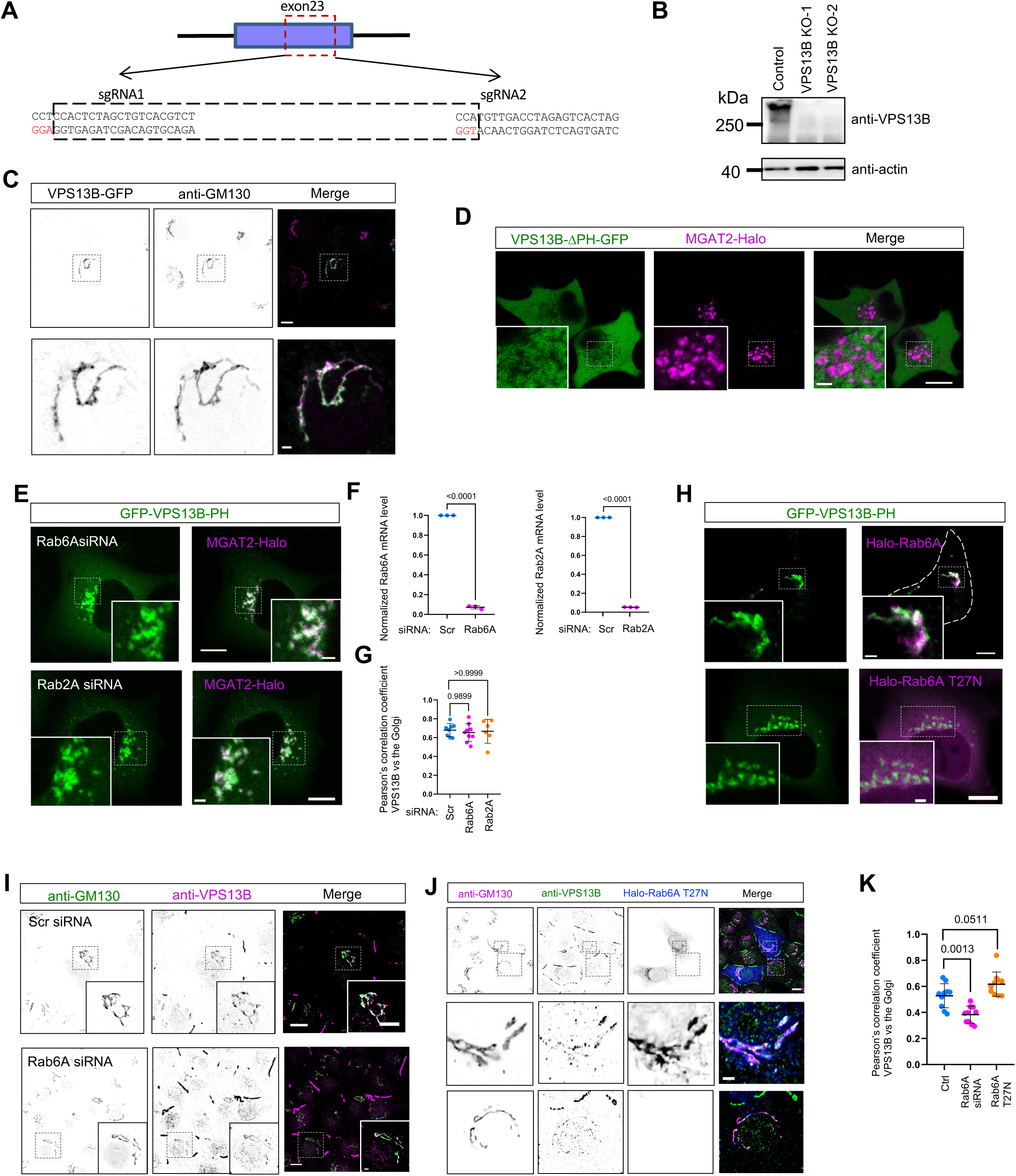
Supplementary data to Fig.1. (A) CRISPR knock-out of VPS13B in HeLa cells (VPS13B-KO). Two sgRNAs are used with the underlined letters, indicating the PAMs (AGG for sgRNA1 and TGG for sgRNA2) for spCas9. (B) Western blots of two VPS13B-KO clones from A. (C) Representative images of fixed HeLa cells transiently expressing VPS13B-GFP (green) stained with GM130 antibody (magenta) with an inset on the bottom. (D) Representative images of live HEK293 cells transiently expressing VPS13B-ΔPH-GFP (green) and MGAT2-Halo (magenta) with an inset on the bottom. (E) Representative images of live HeLa cells transiently expressing GFP-VPS13B-PH (green) along with either Halo-Rab6A (top panel) or Halo-Rab6A T27N (bottom panel) with an inset on the bottom. (F) Representative images of live HeLa cells transiently expressing GFP-VPS13B-PH (green) upon either Rab6A siRNA (top panel) or Rab2 siRNA (bottom panel) treatments with insets. (G) qPCR assays indicated the efficiency of siRNA-mediated suppression of Rab6A and Rab2A from 3 independent experiments as in (**E, F**). Mean ± SD. Two tailed unpaired student t test. (H) Pearson’s correlation coefficient of anti-VPS13B vs the Golgi as shown in (**E, F**) based on at least 10 cells from the three groups in 3 independent experiments. Ordinary one-way ANOVA with Tukey’s multiple comparisons test. Mean ± SD. (I) Representative images of fixed HeLa cells stained with VPS13B antibody (magenta) and GM130 antibody (green) upon either scrambled (top panel) or Rab6A siRNA (bottom panel) treatments with insets. (J) Representative images of fixed HeLa cells expressing Rab6A T27N mutants stained with VPS13B antibody (magenta) and GM130 antibody (green). Two insets were shown on the bottom with or without Rab6A-T27N, respectively. (K) Pearson’s correlation coefficient of anti-VPS13B vs anti-GM130 as shown in (**I, J**) based on at least 10 cells in 3 independent experiments. Ordinary one-way ANOVA with Tukey’s multiple comparisons test. Mean ± SD. Scale bar, 10 μm in the whole cell images and 2 μm in the insets in (C, D, E, F, H& I).

**Fig. S2.**
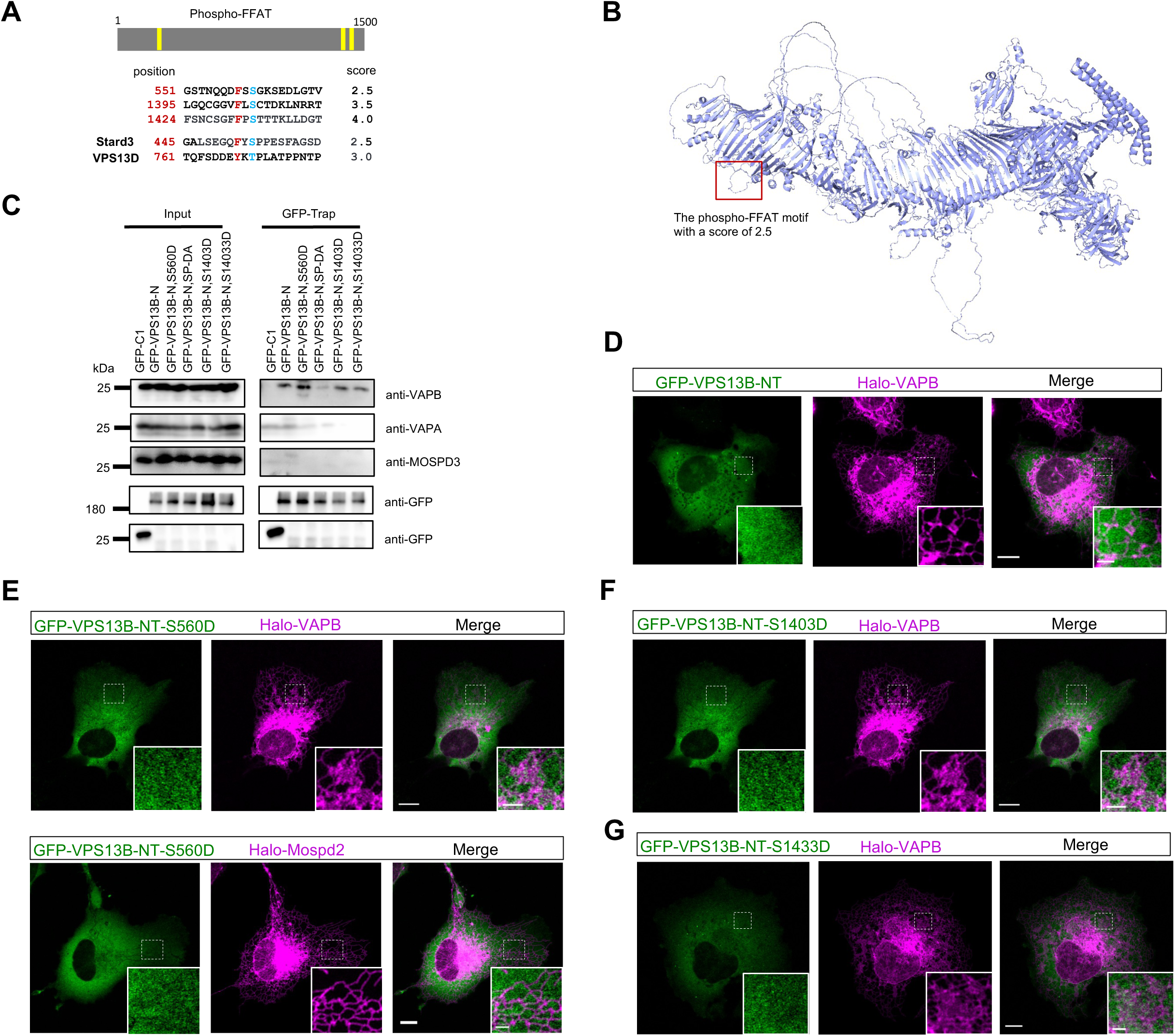
The association between VPS13B and VAPs. (A) Three predicted phospho-FFAT motifs in VPS13B. (B) Alphafold-predicted structures of VPS13B with the predicted phospho-FFAT motif (551-GSTNQQDFSSGKSEDLGTV) highlighted in a red box. (C) coIP assays showed interactions between GFP-VPS13B-NT (1-1500aa) containing different point mutations in phospho-FFAT motifs and endogenous VAPs in HEK293 cells. (D-G) Representative images of HeLa cell expressing either WT or phosphomimic mutants of GFP-VPS13B-NT (green) along with Halo-VAPs (magenta). Scale bar, 10 μm in whole and 2 μm in insets in (D-G).

**Fig. S3.**
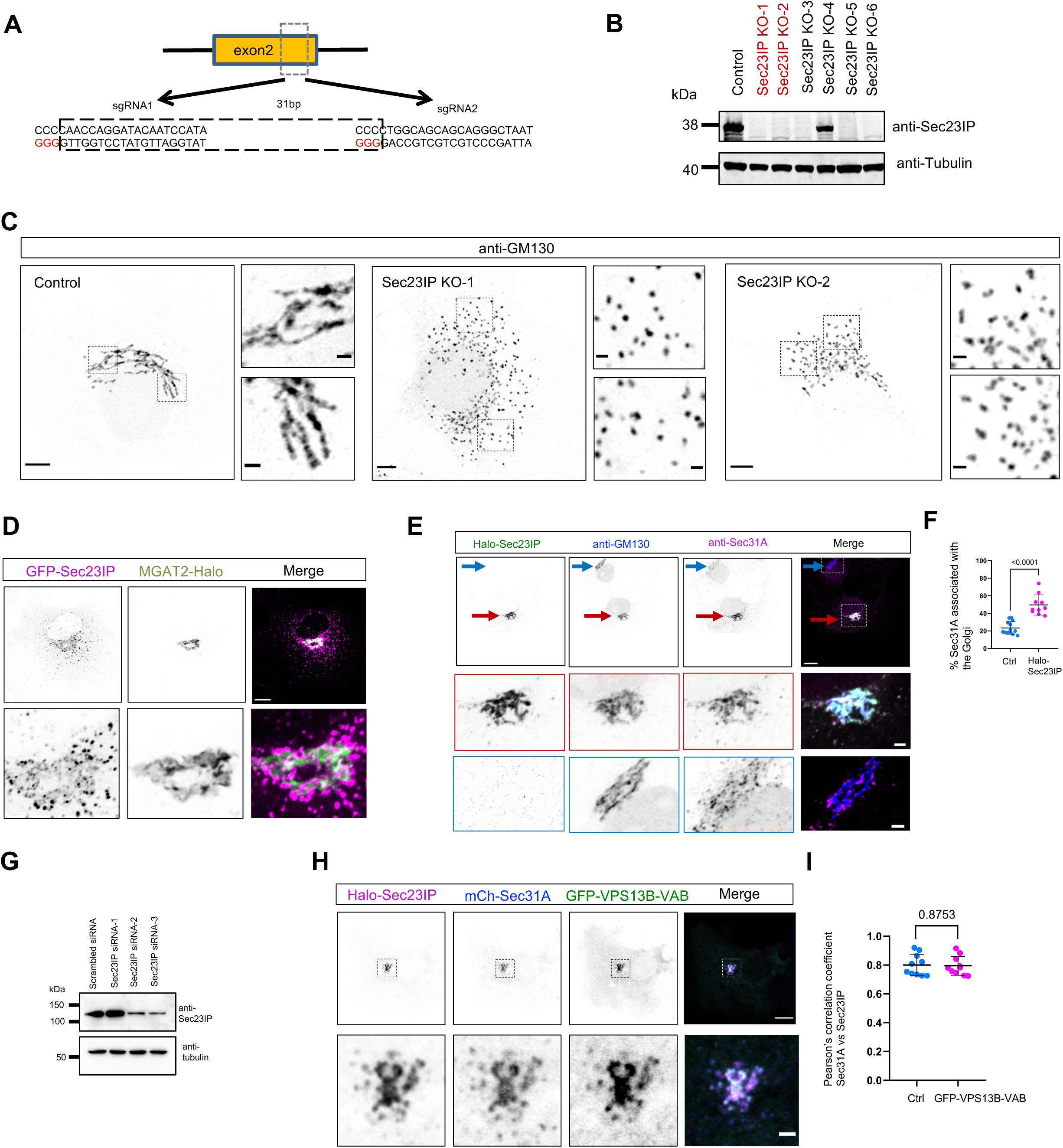
Sec23IP KO results in Golgi fragmentation phenocopying the VPS13B KO in HeLa. (A) CRISPR knock-out of Sec23IP in HeLa cells (Sec23IP-KO). Two sgRNAs are used, indicating the PAMs (GGG for sgRNA1 and sgRNA2) for spCas9. (B) Western blots of six Sec23IP-KO clones in (A). Sec23IP-KO clone-1 and -2 were used in the study. (C) Representative images of fixed control or two Sec23IP-KO clones as in (B) stained with anti-GM130 with two insets on the right. (D) Representative images of a live HeLa cell expressing GFP-Sec23IP (magenta) and MGAT2-Halo (green) with an inset on the bottom. (E) Representative images of fixed HeLa cells expressing Halo-Sec23IP (green) stained with GM130 antibody (blue) and Sec31A antibody (magenta) with two insets on bottom. Red arrows indicated a cell with the expression of Halo-Sec23IP while blue arrows denoted a cell without Halo-Sec23IP expression. (F) Percentage of Sec31A puncta associated with the Golgi in either control (12 cells) or cells expressing Halo-Sec23IP (10 cells) as in (E). Mean ± SD. Two tailed unpaired student t test. (G) Western blots showing the efficiency of siRNA-mediated Sec23IP depletion in VPS13B KO HeLa cells. (H) Representative images of a live HeLa cell transiently expressing GFP-VPS13B-VAB (green), Halo-Sec23IP (magenta) and mCh-Sec23A (blue) with an inset on the bottom. (I) Pearson’s correlation coefficient of Halo-Sec23IP vs mCh-Sec23A as shown in (D) in the absence (10 cells) or presence of GFP-VPS13B-VAB (9 cells) in 3 independent experiments. Mean ± SD. Two tailed unpaired student t test. Scale bar, 10 μm in whole and 2 μm in insets in (C, D, E & H).

**Fig. S4.**
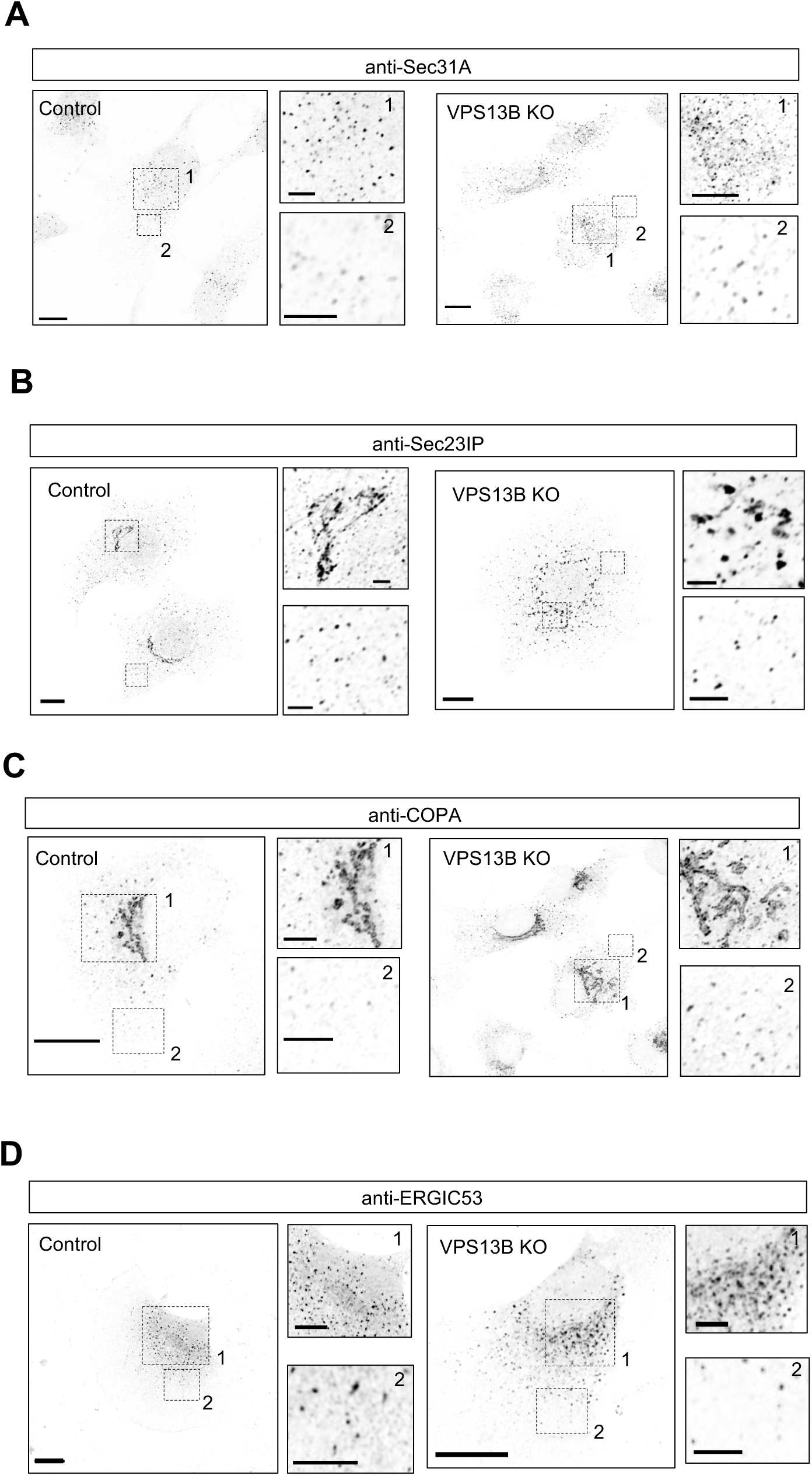
COPII, COPI and conventional ERGIC53 in VPS13B KO HeLa. (A-D) Representative images of fixed control (left panel) or VPS13B KO (right panel) HeLa cells stained with antibodies against Sec31A (A), Sec23IP (B), COPA (C) and ERGIC53 (D) with two insets to the right. Scale bar, 10 μm in whole and 2 μm in insets in (A-D).

**Fig. S5.**
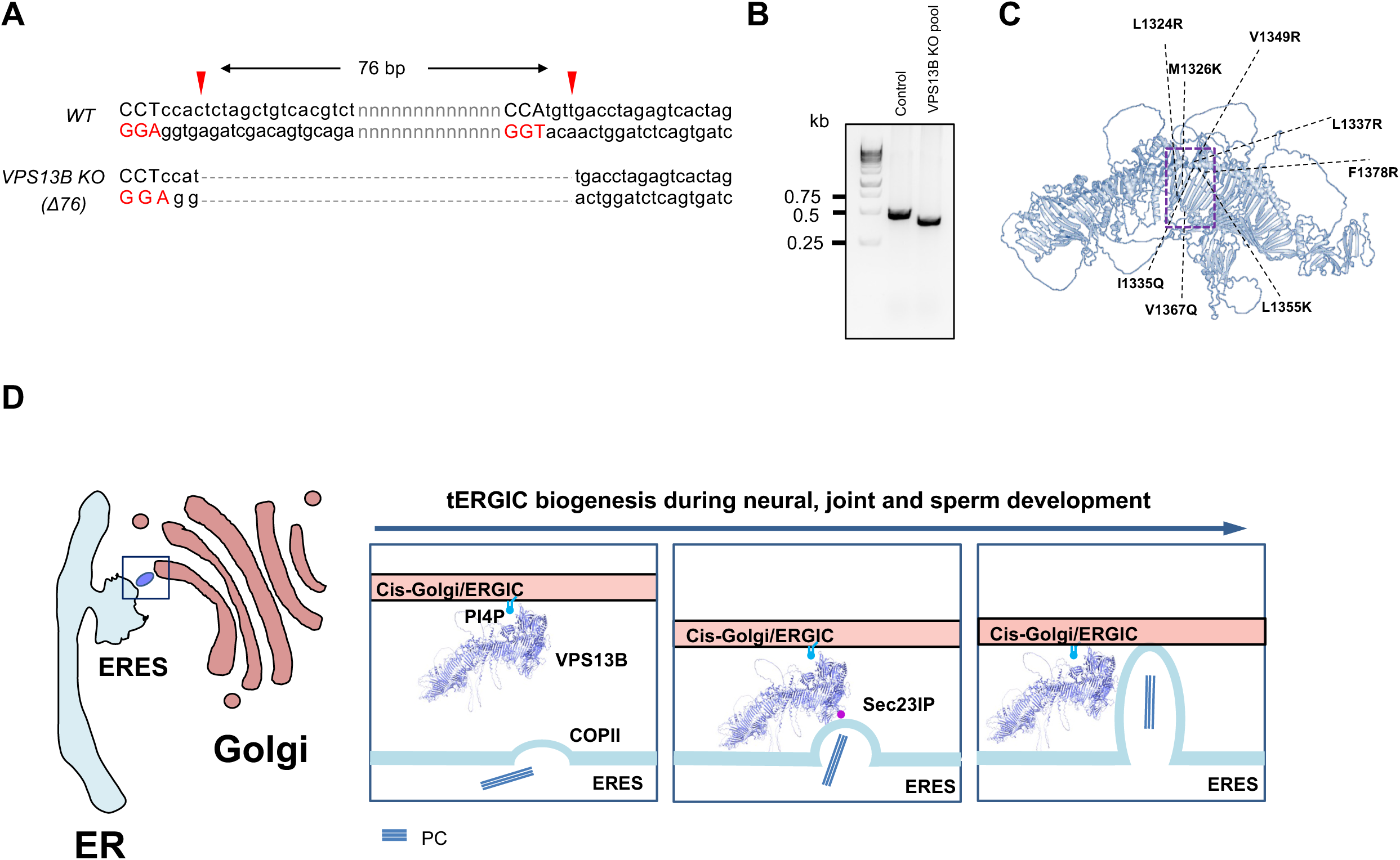
Lipid transfer of VPS13B is required for the growth of tERGIC. (A) CRISPR knock-out of VPS13B in HEK293 cells. Two sgRNAs are used, indicating the PAMs (GGA for sgRNA1 and GGT for sgRNA2) for spCas9. (B) DNA gel of a VPS13B-KO HEK293 pool in (**A**). (C) AlphaFold predicted structures of a VPS13B lipid transfer-defective mutant (VPS13B-LTP^mut^-GFP) with mutated hydrophobic residues in the midway of hydrophobic groove. (D) A working model of VPS13B functions at ERES-Golgi interface. The interaction between VPS13B and Sec23IP promotes the ERES-Golgi association. At ERES-Golgi interfaces, lipid transfer of VPS13B promotes the growth of tERGIC, and thereby meets the requirement for high intensity of proteins/lipids trafficking at certain developmental stages such as the developments of the neural system, joints and sperms. The mode of lipid transfer mediated by VPS13B is under investigation.

## Notes

### Competing Interest Statement

The authors have declared no competing interest.

